# Capturing a crucial ‘disorder-to-order transition’ at the heart of the coronavirus molecular pathology – triggered by highly persistent, interchangeable salt-bridges

**DOI:** 10.1101/2021.12.29.474439

**Authors:** Sourav Roy, Prithwi Ghosh, Abhirup Bandyopadhyay, Sankar Basu

## Abstract

The COVID-19 origin debate has greatly been influenced by Genome comparison studies of late, revealing the seemingly sudden emergence of the Furin-Like Cleavage Site at the S1/S2 junction of the SARS-CoV-2 Spike (FLCS_Spike_) containing its _681_PRRAR_685_ motif, absent in other related respiratory viruses. Being the rate-limiting (i.e., the slowest) step, the host Furin cleavage is instrumental in the abrupt increase in transmissibility in COVID-19, compared to earlier onsets of respiratory viral diseases. In such a context, the current paper entraps a ’disorder-to-order transition’ of the FLCS_Spike_ (concomitant to an entropy arrest) upon binding to Furin. The interaction clearly seems to be optimized for a more efficient proteolytic cleavage in SARS-CoV-2. The study further shows the formation of dynamically interchangeable and persistent networks of salt-bridges at the Spike–Furin interface in SARS-CoV-2 involving the three arginines (R682, R683, R685) of the FLCS_Spike_ with several anionic residues (E230, E236, D259, D264, D306) coming from Furin, strategically distributed around its catalytic triad. Multiplicity and structural degeneracy of plausible salt-bridge network archetypes seems the other key characteristic features of the Spike–Furin binding in SARS-CoV-2 allowing the system to breathe – a trademark of protein disorder transitions. Interestingly, with respect to the homologous interaction in SARS-CoV (2002/2003) taken as a baseline, the Spike–Furin binding events generally in the coronavirus lineage seems to have a preference for ionic bond formation, even with lesser number of cationic residues at their potentially polybasic FLCS_Spike_ patches. The interaction energies are suggestive of a characteristic metastabilities attributed to Spike–Furin interactions generally to the coronavirus lineage – which appears to be favorable for proteolytic cleavages targeted at flexible protein loops. T he current findings not only offer novel mechanistic insights into the coronavirus molecular pathology and evolution but also add substantially to the existing theories of proteolytic cleavages.

## 1. Introduction

There has been a dramatic shift [1] in latest COVID research from its early chapters (2019-20 → 2020-21) brought about by epidemiological and evolutionary (genome comparison) studies of late [2, 3], reporting the presence of an arginine-rich polybasic Furin like cleavage site (FLCS) at the Spike-S1/S2 junction of SARS-CoV-2, absent otherwise in related coronavirus species. This has certainly raised doubts about the origin of SARS-CoV-2 (whether purely natural [4, 5] or otherwise [1, 6]) which is presently obscure and debatable [7, 8]. The latest understanding is that it was during a systematic ’gain of function’ mutational studies [9] carried out on gradually evolving strains of the coronavirus (starting from its natural template, SARS-CoV, 2003) that the virus triggering the current pandemic (SARS-CoV-2) accidentally got released and that too from a highest bio-safety level virology laboratory (at the Wuhan Institute of Virology [10, 11], Wuhan, China). Alarming concerns have thus been unavoidable ever since on the ethical grounds of the decades-long practices of ‘gain of function’ research in virology [12–14]. Apart from the presence of the FLCS in the SARS-CoV-2 Spike (FLCS_Spike_: absent in other beta-CoVs) the other most striking evolutionary feature of SARS-CoV-2 has been its RBD_Spike_ [15] which is highly optimized for binding with its human host receptor, hACE2 (among all species known to harbor homologous receptors [16, 17]), thus channeling the viral influx heavily towards the human population. Both these features, collectively and on their own, present a certain degree of abruptness considering both the time-scale and the sequential nature of changes attributed to natural evolution. These features thereby are strongly argued to carry blueprints of possible human interventions, demanding for further investigations.

The RBD_Spike_ – hACE2 interaction in SARS-CoV-2 had been well characterized by 2019-20 [18–20] and so was its role in the virus host cell entry [18]. A kinetically driven “down-to-up” conformational transition of RBD_Spike_ triggered upon a close proximity to hACE2 has been found instrumental for the virus host cell entry. This “down-to-up” transition of RBD_Spike_ enables it to dock to the solvent exposed hACE2 molecular surface. The RBD_Spike_ – hACE2 interaction, reminiscent of a molecular handshake [15], serves as a molecular switch in the viral cell entry. The RBD_Spike_ – hACE2 interface has a low electrostatic matching [15], characteristic of quasi-stable interactions and therefore perhaps best-fitted for (transient) molecular switches. The befitting surface docked to the solvent exposed Spike binding site of hACE2 selects solely for a standing up conformation of the RBD_Spike_. Interestingly, as revealed by Cryo-EM studies [21] in SARS-CoV, this ’up’ state is the temporally prevalent state while in SARS-CoV-2, the ’up’ state is only triggered in close proximity to hACE2. This, when viewed together with the abrupt emergence of host Furin cleavage and that of the polybasic FLCS_Spike_ site in SARS-CoV-2 (absent in SARS-CoV) certainly appears to be contradictory to the slow and graduated changes attributed to natural evolution. The more efficient the cleavage, the more easier it would be for the virus to gain entry inside the host cell [1]. At one end, there’s no FLCS_Spike_ in the SARS-CoV Spike which therefore is only cleaved randomly (and hence less frequently) by non-specific proteases, thereby always demands an ’up’ state of the RBD_Spike_ for the interaction to occur [22, 23]. At the other end, the native, resting, lying down state of the RBD_Spike_ in SARS-CoV-2 helps the virus to escape the host immune surveillance mechanisms [24, 25]. The overall impact is interesting (and perhaps somewhat counter-intuitive) that there is an increase in binding affinity in the younger homologue (SARS-CoV-2) compared to its evolutionary ancestor (in SARS-CoV, 2002/2003) while in terms of binding stability (attributed to electrostatic matching at the interface) there is a ’critical’ drop [26] from CoV to CoV-2, imparting a bouncing nature in the later interaction and thereby making the neighboring cells more susceptible to subsequent viral entries than the earlier event (SARS-CoV).

In addition to the RBD_Spike_, host protease pre-activation (or priming) plays an imperative role in SARS-CoV-2 pathogenesis critically enhancing its efficient host cell entry. Notably, the priming step acts a promotional factor in SARS-CoV-2 but is non-essential (for infection and cell-cell fusions) [27] generally in the coronavirus lineage. In case of SARS-CoV-2 (and, its subsequent lately evolved variants [28–30]) the virus uses its arginine-rich FLCS_Spike_ as a lucrative bait to recruit host-encoded pro-protein convertases (PC), primarily Furin [31] (the best-characterized mammalian PC [32]) rapidly at the Spike–S1/S2 junction. This then is relatively slowly [1] followed by the host Furin cleavage of the said junction (Spike–S1/S2) eventually leading to a more efficient host cell entry of SARS-CoV-2 [33] (compared to SARS-CoV and other related respiratory viruses) and its variants [1,28,30]. Apart from Furin, SARS-CoV-2 entry is also primed by a cell surface protease TMPRSS2, lysosomal cathepsins [34], and, also by proteases (NE, PR3, CatG, NSP4) released by activated neutrophils [35] swarming around the invaded pathogen to elicit an immune response. They therefore can be hijacked by virus-derived surface proteins as a mean to escape the host immune surveillance. All these eventually leads to a cumulative effect of (Furin like) host proteases on SARS-CoV-2 entry. Furin is also well known for its undue involvement in various pathologies, especially related to bacterial and other viral diseases (e.g., Anthrax, Ebola) [32]. Structurally, it is the association of two domains, (i) a catalytic domain consisting of closely packed α-helices and intertwined crisscrossed beta strands at the N terminus and (ii) an all-β (C-terminus) P domain. While the evolutionarily varied domain is the P domain, the catalytic domain is highly conserved across mammals, and, further harbors a characteristic ‘Serine–Histidine–Aspartate’ catalytic triad that mediates the (proteolytic) cleavage. With the help of this triad, Furin cleaves precursor proteins (or, pro-proteins) having a basic consensus of ‘R-X-K/R-R-’ like motif with stringent specificity [32]. The stringency, as well as the preference towards basic residues, has been structurally explained in Furin by the presence of contoured surface loops shaping the active site and harboring highly charge-complementary pockets. The FLCS_Spike_ in SARS-CoV-2 that recruits Furin with high affinity has a ‘*_681_PRRAR_685_*’ motif as the polybasic consensus, encoded by a 15-nucleotide insert 5′-CCTCGGCGGGCACGT-3′ at the Spike–S1/S2 junction. Interestingly, the two consecutive arginine residues (R) in the ‘PRRAR’ are both encoded by a CGG codon (2^nd^, 3^rd^ triplet in the ORF^1^) - which is the most infrequent codon for viruses (out of the six codons dedicated for arginines) [1]. The presence of this unique and specific 15-nucleotide sequence in the SARS-CoV-2 genome was even satirically referred to as a “smoking gun” [1] supporting the lab-origin theory of this virus. Further, as confirmed by functional studies, the loss of FLCS_Spike_ has been shown to attenuate SARS-CoV-2 pathogenesis [33]. The discovery of the unique presence of FLCS_Spike_ in SARS-CoV-2 has also caused a shift of scientific parlance in the subject, directing researchers from a physics-to a chemistry-observation window. The RBD_Spike_–hACE2 interaction in SARS-CoV-2 is essentially (bio-)physical, guided by the physical laws of diffusion and collision, applicable to non-covalent (*van der Waals*) inter-atomic interactions. While, the next important step involves breaking of a covalent bond in the FLCS_Spike_ by Furin and therefore is a chemical process [1]. Naturally, the later is much slower than the earlier, further making it (FLCS_Spike_–Furin) the rate-limiting step in the viral host cell entry [1]. Insertion of the polybasic activation loop (_681_PRRAR_685_) in the SARS-CoV-2 Spike as opposed to other related respiratory viruses has been pinned to increased virulence, transmissibility, and pathogenesis [35–37]. This makes it critical to understand in tandem the dynamics and transition of these loops into ordered conformers upon binding to Furin (or other proteases) to pinpoint the molecular interactions in play. Such knowledge could be pivotal in expanding our understanding of how loop dynamics, interaction and stabilization forms the molecular basis of enhanced pathogenicity and virulence of SARS-CoV-2 Spike glycoproteins. Most structures of Spike glycoproteins solved till date are in their pre-fusion state – which is its active state, relevant for both the FLCS_Spike_–Furin and the RBD_Spike_–hACE2 interactions. The pre-fusion forms, aimed to address different specific research queries related to the coronavirus molecular pathology, are either mutated or abrogated at the FLCS_Spike_ region (i.e., the Spike–S1/S2 junction) or otherwise engineered with stabilizing mutations [21,24,38], aimed at obtaining overall structural information. However, even with these stabilizing modulations, all cryo-EM coronavirus Spike structures lack experimental (primary) data, and, hence, atomic coordinates for their FLCS_Spike_ patches (missing stretches of 10-12 amino acids [39]) harboring the polybasic activation loop (*_681_PRRAR_685_*) in SARS-CoV-2 Spike, and, equivalent homologous pentapeptide sequence motifs in other coronavirus Spikes. The pre-fusion structure of SARS-CoV-2 Spike (PDB^2^ ID: 6XR8) [21] could decipher the structure of 25 ordered peptides downstream of S2 hitherto unreported including residues of N terminus, several peripheral loops and glycans [21]. Even this most insightful high resolution structural study, which could reveal the distinct conformational states (pre- and post-fusion forms) of the SARS-CoV-2 Spike, was unable to ascertain the conformation of the surface exposed disordered FLCS_Spike_ loop harboring the Spike–S1/S2 junction [21]. The region is further clearly and categorically declared as a *“surface exposed disordered loop”* [21]. The presence of such ‘disordered activation loops’ (also called ‘natively unstructured loops’ [40] or, less technically, ‘flexible’ loops/regions) at proteolytic sites is common to many protease families (e.g., Caspases, Furins, Rho-activated kinases etc.) and in spite of decades long controversies are still assumed to serve as key structural and kinetic determinants of protease substrates [40]. The loops, often making the substrates more susceptible to protease binding and cleavage [31], proteasomal degradation [41], phospho-regulation and consequent priming of viral pathogenesis [42] are believed to have been evolved often from sensitive and fragile globular-disorder intermediates like coiled-coil assemblies [43]. Having said that, there is little experimental structural information that is available on the cleavage loops for our current subjects, the coronavirus Spikes.

The loop-disorder is further supported by extensive (coarse-grained) multi-microsecond molecular dynamic (MD) simulations of the representative SARS-CoV-2 Spike structures (6VXX, closed state; 6VYB: partially open state, ectodomain) [36] with modeled FLCS_Spike_ loops (residue ranges: 676-690) [44] that show violation of ergodic hypothesis [36], hinting towards a flexible albeit biased conformation of the activation loop for its function. In other words, during the entire course of these long MD simulations, the loops remained largely unstructured [36] sampling several outwardly extended conformations making them attractive and accessible cleavage sites for Furin and other pro-protein convertases. So the loop disorder does not appear to be short-termed, rather dynamically sustained in the free form of the Spike. As an alternative and arguably a complementary approach, the same paper further models an ensemble of loops *ab initio* [36] via RosettaRemodel [45] from the closed state Spike structure (6VXX). Subsequent to the modeling, followed by clustering and sorting based on energy, the lowest energy conformation was retained and further refined by kinetic closure [46]. In the conclusive remarks [36], the authors comment that the *ab initio* modeling indicated that there might be formation of short helices near the cleavage site of the loop; however, these observations are not really supported by any data and/or geometric analysis such as the Ramachandran Plot. The very fact that the FLCS_Spike_ (in its *_681_PRRAR_685_*) contains pockets of heavily localized positive charge cloud (a common cause of protein disorder [47]) makes it naturally and intrinsically prone to structural disorder. This makes the analysis even more interesting and non-trivial, and further implies that there needs to be a ‘*disorder-to-order transition*’ [47] of the said loop (i.e., the FLCS_Spike_ region) upon interaction (or binding) with Furin. That is because the binding needs to be strong enough to sustain the disordered substrate (FLCS_Spike_) jammed into metastable intermediate conformation(s) that support the efficient cleavage of the Spike–S1/S2 junction by the Furin catalytic triad. Recent experimental studies primarily based on functional assays have referred to the ‘ostensibly unresolved’ FLCS_Spike_ as a ‘protruding out loop-like structure’ in the exterior [33] of the SARS-CoV-2 Spike, away from the RBD_Spike_. Others [48] have completely neglected the plausible conformational variation (entropy) of the missing patch(es) in their molecular docking and dynamics studies. Thus, little has genuinely been explored structurally on the Spike–Furin interaction. Given this background, here, in this paper, we present a rigorous structural dynamics study with strong theoretical rationales to penetrate deeper into the Spike–Furin interaction in SARS-CoV-2 in an atomistic detail. To the best of our knowledge, we are the first group to report on the key ‘*disorder-to-order* transition’ occurring at the SARS-CoV-2 Spike–Furin interface. Based on present knowledge and understanding, the revealed transition together with the disorder, intrinsic to the CoV-2 FLCS_Spike_ [21] should find its place at the very heart of the coronavirus molecular pathology. We further demonstrate that the involved ‘*disorder-to-order* transition’ in CoV-2 is triggered and sustained by highly persistent interchangeable salt-bridge dynamics – which is often characteristic to protein disorder transitions [49, 50]. We also present a novel application of the legendary Ramachandran Plot [51] in probing the said ‘transition’ at the CoV-2 Spike–Furin interface. The conclusions should also hold true for lately evolving CoV-2 variants [28, 30].

## 2. Materials and Methods

### 2.1.1. Details and rationales of experimental structural templates used for docking and dynamics

Following earlier studies on the use of the SARS-CoV-2 Spike glycoprotein [15,52,53], the cryo-EM structure of its pre-fusion form (PDB ID: 6XR8, 2.9 Å, 25995 protein atoms [21]) was used as its representative structure (receptor). For the host protease, Furin (ligand), we first made a thorough structural survey of its domains and enzyme active sites, especially that of the catalytic domain and triad). There are only a few experimental (X-ray) structures of Furin to be found presently in the Protein Data Bank [39] (PDB ID: 1P8J, 5JXH) which are of equivalent resolutions with insignificant structural deviations (C^α^-RMSD: 0.31 Å) upon alignment. More importantly, the region spanning Catalytic residues in both are well conserved. 1P8J, (resolution: 2.6 Å), the first Furin structure ever to be solved for Furin [32] is undoubtedly the most studied and best characterized Furin structure to have, also insightful in terms of proteolytic cleavage mechanisms [48,54–57]. It is for these reasons, 1P8J was chosen as the representative host protease (Furin) structure in the current study. Again, 1P8J has 8 identical chains (pairwise RMSD < 0.2 Å) in its asymmetric unit combining to as many as 11 bio-assemblies. The majority of the bio-assemblies (bio-assemblies: 3-10) are monomeric – which is the interactive molecular species (or, functionally effective bio-assembly) involved in the Spike–Furin interaction [48]. This made us further retain the first chain (chain A) of 1P8J alone.

Further, to set an appropriate baseline for the Spike–Furin interaction in SARS-CoV-2, a second round of docking was performed by taking the same ligand (Furin: 1P8J) and docking it onto the representative homologous Spike structure (PDB ID: 7AKJ) [58] of SARS-CoV (2002/2003). This cryo EM structure was solved in the process of exploring ‘the convalescent patient sera option’ for the then seemed possible prevention and treatment of COVID-19, as a natural extension of SARS [59]. The neutralizing antibody Fab fragments (chains: D-H & L in 7AKJ), used to pull down the SARS-CoV Spike, binds at the top of the Spike canopy or the Spike S1 subunit involved in receptor binding [58] – which is situated way too far from the Spike–S1/S2 interface (harboring its FLCS_Spike_) to have any realistic interference with the Spike–Furin binding. The Spike–Furin guided docking in SARS-CoV (to be discussed in section **2.3.1.3**) was followed by repeating all subsequent baseline structural dynamics calculations for the Spike– Furin complex. To set this appropriate baseline is indispensable particularly for the quantitative interpretation of equilibrium thermodynamic parameters of the Spike–Furin binding, due to be discussed in later sections (section **2.6**).

### 2.1.2. Dataset of coronavirus Spikes

For an evolutionary analysis of sequence based disorder predictions, a second dataset was compiled consisting of 44 experimentally solved (exclusively cryo-EM) structures (**Table S1, Supplementary Materials**) of the pre-fusion form CoV/CoV-2 Spike culled at a resolution of ‘not worse than 3 Å’ from the PDB. These coronavirus Spike structures are solved to serve different specific research objectives at various pH and other varying physico-chemical conditions [17,58,60,61]. All these structures were found not to have any experimental (primary) data for its FLCS_Spike_ patch (located at their Spike–S1/S2 junctions) resulting in missing atomic coordinates (remarked under ‘REMARK 465’ in the corresponding PDB files) for the said patch (10-12 amino acids).

### 2.2. Modeling of missing disordered loops in Spike

To explore the interaction dynamics between SARS-CoV-2 Spike (PDB ID: 6XR8) and Furin (1P8J), first we needed to have ‘all-atom’ atomic models for both partners. Furin (1P8J) is much smaller (468 residues) compared to the Spike homo-trimer (3 identical chains with 1149 residues per chain) and has no missing stretches in its X-ray structure but for the terminal most residues. The Spike trimeric structure, however, being a large glycoprotein, characteristically contains missing stretches of residues (of roughly similar lengths) localized at strategic positions, adding up to missing coordinates for 42 amino acids in 6XR8. As discussed in the previous subsection (section **2.1.2**), to have such missing stretches of residues, especially at the Spike–S1/ S2 junctions, is common to all Spikes (**Table S1, Supplementary Materials**) irrespective of the particular coronavirus species. These missing stretches map to flexible [36] as well as disordered [21] loop protrusions resulting from the extended internal packing involved in the trimerization of the monomeric S units. The missing disordered stretches in the representative SARS-CoV-2 Spike pre-fusion structure (6XR8) were then modeled by MODELLER (v.10.1) [62] using its full-length (proteomic [63]) sequence (https://www.rcsb.org/fasta/entry/6XR8/display) obtained from the Protein Data Bank [39]. To account for the loop disorder in the modeled missing stretches (in 6XR8), an ensemble modeling approach was adapted. To that end, the ‘*automodel*’ module of MODELLER was implemented in an iterative cycle of 500 runs, producing that many conformationally non-redundant ‘all-atom’ trimeric Spike atomic models, only varying among themselves at the modeled missing stretches.

Missing residues for the baseline structure of SARS-CoV Spike (7AKJ) were also build in a similar fashion using MODELLER, though opting for a much reduced space of conformational sampling (50 runs of ‘*automodel*’). The lowest energy model among these (ranked by the all-atom Rosetta energy function, availed through *<u>Rosetta@home</u>* [64]) was retained as the representative ‘all-atom’ SARS-CoV Spike structure to be used for all subsequent baseline calculations. This sampling may be considered adequate to represent the mildly varying (reduced) conformational space of the missing FLCS_Spike_ patch (_664_SLLRSTS_670_, https://www.rcsb.org/fasta/entry/7AKJ/display) in 7AKJ – which is much shorter than its homologous missing patch in 6XR8 (_677_QTNS**PRRAR**SVA_689_). The FLCS_Spike_ in SARS-CoV is further composed mostly of either small-polar (serines, threonines) or hydrophobic (leucines) side chains with constrained rotameric variations. Noticeably, the single charged residue (R667) situated midst the 7 residue missing patch in 7AKJ is conserved (as R685) in its evolutionarily descendant sequence in 6XR8 (**Figure S1, Supplementary Materials**).

### 2.3.1. The Spike–Furin Molecular Docking Simulations

#### 2.3.1.1. Blind ab-initio docking in Cluspro 2.0

Subsequent to filling up for the structural voids in the trimeric SARS-CoV-2 Spike (section **2.2**), an ensemble docking approach was adapted (using ‘blind docking’ in ClusPro 2.0 [65]) wherein Furin (PDB ID: 1P8J, ligand) was docked *ab initio* onto each unbound ‘all atom’ Spike atomic model (receptor) belonging to the disordered FLCS_Spike_ ensemble. The ClusPro web interface performs *ab initio* blind docking in successive steps, combining course grain sampling and fine grain refinements. First, a grid-based rigid-body docking is performed between the receptor (static, fixed at the center of a cubic box) and the ligand (dynamic, placed on a movable grid) in PIPER [66] using its fast Fourier transform (FFT) correlation approach, sampling billions of conformations. The PIPER computed interaction energies (for poses sampled at each grid point) are produced in the form of correlation functions that make the scoring compatible with FFTs, rendering high numerical efficiency of sampling. The latest PIPER–derived scoring function is an upgraded variant of the original internal energy function (https://www.vajdalab.org/protein-protein-docking) which is based on the sum of terms representing shape complementarity, electrostatic, and desolvation contributions. A large number of unrealistic poses are then filtered out by PIPER and the 1,000 lowest-energy poses are retained. These are then undertaken to RMSD^3^–based structural clustering (using a relaxed 10 Å cutoff for iRMSD [67]), wherein, largest clusters [65] are retained, representing the most likely poses. While the largest clusters do not necessarily contain the most near-native poses, the success rate is generally quite high across families of protein complexes [65]. Usually ten or fewer clusters (∼10-12 pose/cluster) are retained at this stage adding up to 100-120 docked poses per run. The final refinement is then performed on these selected poses by rigorous rounds of energy minimization. The blind ‘ensemble docking’ thus performed between Furin (ligand) and each Spike model (receptor) does not impose any additional active-site or contact residue constraints. This resultant initial pool of Spike–Furin docked poses under each docked ensemble (i.e., for each Spike model) had in them the ligand docked onto widely varying sites spread all over the trimeric Spike receptor – which were then pulled down into one unsorted set. Since the ‘desired interface’ would necessarily involve the FLCS_Spike_, the pentapeptide motif (_681_PRRAR_685_) was then used as ‘contact residue filters’ to discard obviously and/or trivially incorrect docked poses. To that end, buried surface areas (see section **2.3.2.1**) and shape complementarities (see section **2.3.2.2**) estimated at the desired docked interfaces were used to filter the ‘plausible’ docked poses in two successive rounds. These filtered ‘plausible’ poses were then further re-ranked by a carefully designed scoring function (see section **2.3.2.1**) optimally combining the steric effects of the two high-level structural descriptors. Thus, in effect, the blind docking could be made to work in a guided manner. The top ranked docked pose (RR1_CoV-2_) thus obtained, was further taken into long MD simulations (see section **2.4**) and subsequent analyses of its structural dynamics.

#### 2.3.1.2. Cross-validation by guided docking in ‘Zdock + IRaPPA re-ranking’

The cluspro – docking was further cross-validated by Zdock with its combined feature of IRAppA re-ranking [68] implemented. To that end, the Spike chains (receptors) were extracted from the top (re-)ranked (see section **2.3.1.1**) cluspro – docked pose, RR1_CoV-2_, and, Furin (1P8J, ligand) was docked onto it with the three arginines (R682, R683, R685, from the first of the three symmetry-related identical Spike chains) pertaining to the _681_PRRAR_685_ motif (in FLCS_Spike_) specified as contact residues on the receptor molecule. No contact residues were specified from the ligand molecule. The returned top ranked model (ZR1_CoV-2_) was retained and further simulated (see section **2.4**) for all subsequent structural dynamics analyses.

#### 2.3.1.3. Setting up appropriate baselines: The SARS-CoV Spike–Furin guided docking in ‘Zdock + IRaPPA re-ranking’

In continuation to the earlier discussion (in section **2.1.1**) on setting up appropriate baselines to interpret the Spike–Furin interaction in SARS-CoV-2, Furin (1P8J, ligand) was docked onto the SARS-CoV Spike (7AKJ, receptor) in Zdock (+IRaPPA re-ranking) in yet another independent docking exercise. To that end, a direct guided mode of docking (similar to ZR1_CoV-2_) was adapted, wherein the whole FLCS_Spike_ patch (residues originally missing in the experimental structure of 7AKJ) was specified as plausible contact residues on the Spike (receptor). This missing FLCS_Spike_ patch build by MODELLER (see section **2.2**) maps to residues 664-670 (_664_SLLRSTS_670_) and those pertaining to the first of the three symmetry-related identical Spike chains were specified as the Spike contact residues for Furin. Again, (in the same spirit to that of ZR1_CoV-2_) no contact residues were specified from Furin. The returned top ranked model (ZR1_CoV_) was retained, simulated (see section **2.4**) and used as a baseline for all subsequent structural dynamics calculations, probing for the absence of the ‘PRRAR’ motif in FLCS_Spike_.

### 2.3.2. Docking scoring, ranking and re-ranking

#### 2.3.2.1. Buried Surface Area calculations

For initial filtering of (Spike–Furin) docked poses (ClusPro 2.0) the atomic accessible surface areas (ASA) were calculated by NACCESS [69] traditionally following the Lee and Richards algorithm [70] for all heavy atoms pertaining to each partner protein molecule (Furin and Spike) in their bound and unbound forms. For the unbound form, the two partner molecules in the bound docked-pose were artificially physically separated and each of them was considered independently, in isolation. The atomic accessibilities were then summed up for their source residues. For each (i^th^) residue belonging to a docked pose, two ASA(i) values were obtained, one for its bound form (ASA_bound_(i)) and the other for its unbound form (ASA_unbound_(i)). A residue falling in the interface would thus have a net non-zero change (ΔASA(i)≠0) in its two ASA(i) values. In other words, these would be the residues at the interface which gets buried upon complexation and are characterized by a net non-zero buried surface area (BSA(i)>0).

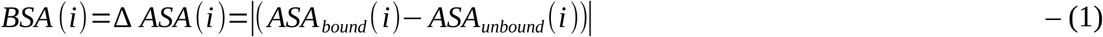

For the case of Spike–Furin docking (SARS-CoV-2), BSA was used as an initial filter to select those interfaces alone that harbors the FLCS_Spike_ loop. In the process, the obviously incorrect poses with Furin docked elsewhere in the Spike were discarded. This was achieved by monitoring BSA_PRRAR_, the summed up BSA for the residues belonging to the pentapeptide _681_PRRAR_685_ motif. The poses that have BSA_PRRAR_>0 were then filtered and retained from the initial pool of *ab-initio* docked poses.

BSA_PRRAR_ for this BSA – filtered set of docked poses was further normalized by the total ΔASA (summed over all residues pertaining to the docked pose) to render the normalized buried surface area for the docked pentapeptide surface patch (nBSA_PRRAR_). The normalization can be expressed by the following equation.

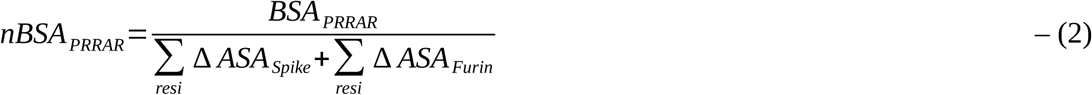

where, although, ‘resi’ stands for all residues pertaining to the protein chain (‘subscripted’ to the corresponding ΔASA sum-over term), it is the interfacial residues (ΔASA(i)≠0) that alone actually contribute to the denominator. The use of nBSA defined at protein-protein interfaces [71] can be considered analogous to that of the surface overlap parameter [72, 73] which has been used extensively in tandem with shape complementarity to study packing within protein interiors.

Wherever applicable, burial (bur) of solvent accessibility for a protein residue (X) was computed (following standard methods [72–75]) by taking the ratio of its ASA when embedded in the protein to that when in a Gly-X-Gly tripeptide fragment with its fully extended conformation.

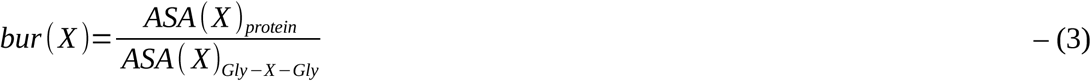

Standard binning techniques for residues based on burial [72–75] were then adapted, with an ever-so-slight modification (opting for 3 instead of 4 burial bins) based on the current requirement. A protein residue based on its burial (defined in the range of [0,1]) could thus be classified into one of the three ‘burial’ classes: (a) buried (*0.0 ≤ bur ≤ 0.05*), (b) partially exposed (*0.05 < bur ≤ 0.30*) or (c) exposed (*bur > 0.30*).

The analysis of residue-wise burial was particularly intended to survey the Furin structure, in order to access its intrinsic propensity to bind to FLCS_Spike_ like disordered and/or flexible loops that harbors patches of highly localized and dense positive charge clouds (e.g., the ‘PRRAR’ pentapeptide motif).

#### 2.3.2.2. Shape Complementarity

For a chosen docked pose which has passed the initial BSA filter (section **2.3.2.1**), the shape complementarity [76, 77] at its interface was computed by the shape correlation (Sc) statistic originally proposed, formulated and programmed (as the sc program, part of the CCP4 package [78]) by Lawrence and Colman [76]. Sc is a correlation function defined in the range of -1 (perfect anti-complementarity) to 1 (perfect complementarity). It elegantly combines both the alignment as well as the proximity of interacting surfaces and is essentially local in nature (resulting from Van der Wall’s packing). Higher the Sc, better the packing at the interface. Well-packed protein-protein interfaces (irrespective of their biological origin and size) usually hit a thin optimal range of Sc values (∼0.55-0.75) [15, 76].

For the case of Spike–Furin docking (SARS-CoV-2), Sc was computed for the BSA – filtered (see section **2.3.2.1**) interfaces alone which refers to those docked poses that harbors the FLCS_Spike_ loop in its interface. Furthermore, for Sc, we narrowed down our observation window to the docked FLCS_Spike_ loop alone – which was necessary and sufficient for the cause of scoring and re-ranking the initially selected (plausible) docked poses. To that end, Sc of the corresponding surface patch (Sc^FLCS^) was computed against its docked surface patches (combined) coming from Furin. Sc is local in nature and can be directly computed for and segregated among pairs of interacting surface patches in multi-body interactions [72, 73]. To render an accurate Sc^FLCS^ for each chosen docked pose, the sc (CCP4) [78] – input file was made to retain coordinates of the corresponding docked patch (resi. 677-688) alone for the interacting Spike chain, appended with coordinates for all atoms pertaining to the docked Furin molecule.

#### 2.3.2.3. The S_dock_ score

The S_dock_ score, designed for the purpose of re-ranking of BSA – filtered (plausible) *ab-initio* docked poses was computed by the following equation.

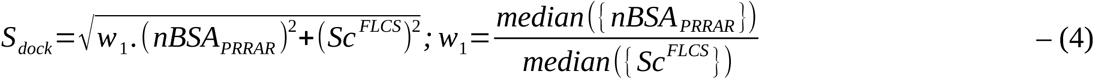

- where, w_1_ (=5.78) appropriately weighted the two components (nBSA, Sc) of the score. In effect, the S_dock_ score elegantly combined the surface fit and overlap at the Spike–Furin interface. The BSA – filtered (plausible) poses were then scored and re-ranked by S_dock_. The design of S_dock_ can be considered analogous to deriving the magnitude of the resultant of two mutually orthogonal vectors in 2D Euclidean space.

### 2.4. Molecular Dynamic Simulations

The top ranked docked SARS-CoV-2 Spike – human Furin complex (RR1_CoV-2_) consisting of 60224 atoms (inclusive of Hydrogens) were used as the initial structural template for an explicit water all atom Molecular Dynamics (MD) simulation run. All MD simulations were performed using Gromacs v2021.3 [79] with OPLS-AA force field [80]. Force-field parameters for the surface-bound glycans of SARS-CoV-2 Spike (6XR8) were used directly from a recent earlier study on the same protein from this laboratory [26], while those for SARS-CoV Spike (7AKJ) were built in an identical manner using the glycoprotein builder at GLYCAM-Web (www.glycam.org) [81]. Cubic box of edge dimension 225.1 Å, solvated by a total (N_sol_) of 348608 TIP3P water molecules was used to solvate the protein complex with an application of periodic boundary conditions of 10 Å from the edge of the box. The system was then charge-neutralized (N_ion_) with 21 Na^+^ ions by replacing the TIP3P waters. The size of the hydrated system thus amounted to 899539 (protein+non-protein) atoms. Bond lengths were constrained by LINCS algorithm [82] and all long-range electrostatic interactions were determined using the smooth particle mesh Ewald (PME) method [83]. Energy minimization was performed with steepest descent algorithm until convergence (∼1000 steps) with a maximum number of steps set to 50000. All simulations were performed at 300K. Temperature equilibration was conducted by the isochoric-isothermal NVT ensemble (constant number of particles, volume, and temperature) with a Berendsen thermostat [84] for 100 ps. The system was then subjected to pressure equilibration in the NPT ensemble (constant number of particles, pressure, and temperature) for 100 ps using the Parrinello–Rahman protocol [85], maintaining a pressure of 1 bar. Coordinates were written subsequent to necessary corrections for Periodic Boundary Conditions (PBC) using the GROMACS command ‘gmx trjconv’ using its -pbc option. Backbone RMSDs were computed using the GROMACS command ‘gmx rms’ and monitored throughout the trajectory. For RR1_CoV-2_, the production run was set to 300 ns which may be considered sufficiently long as the system showed convergence. For cross-validation purposes, a second MD simulation was set up with ZR1_CoV-2_ as the template (see section **2.3.1.2**) and run for 100 ns with identical cubic box dimensions, N_sol_ and N_ion_. To set up appropriate baselines, yet another third independent simulation was set up and run for 100 ns using ZR1_CoV_ as the template (see section. **2.3.1.3**) with cubic box of edge dimension 190.6 Å, solvated by 214418 water molecules and charge-neutralized by 45 Na^+^ ions. All three simulation trajectories attained equilibrium, ensuring seamless downstream calculations. For subsequent analyses of structural dynamics, structural snapshots were extracted at a regular interval of 10 ps from all three trajectories leading to 30000 snapshots for RR1_CoV-2_ (the subject), 10000 snapshots for ZR1_CoV-2_ (the subject for cross-validation) as well as ZR1_CoV_ (the baseline). All simulations were performed on a local workstation with Gromacs v2021.3 [79] with CUDA acceleration v11.2 powered by an NVIDIA RTX 3080 GPU with 8704 CUDA compute cores resulting in an average output simulation trajectory of ∼ 8.4 ns/day.

### 2.5.1. Identifying Salt-bridges at the Spike–Furin interface

For each selected Spike–Furin interface which could either belong to the static ensemble of top ranked docked poses (see section **2.3.2**) or time-frames/snapshots pertaining to MD simulation trajectories (see section **2.4**) produced from top ranked selected docked poses, first, its interfacial inter-residue contact map was extracted. An interfacial inter-residue contact at the said interface was defined and detected when two heavy atoms, one coming from a Spike– and one coming from a Furin– residue was found within 4.0 Å of each-other.

Further, from this interfacial contact map, ionic bonds / salt-bridges were identified following standard definitions and computational techniques [49,50,86,87]. To that end, those inter-residue contacts were assembled and characterized as salt-bridges / ionic bonds where two oppositely charged side chain heavy atoms, a nitrogen (N^+^) and an oxygen (O^-^), coming from two different amino acid residues from the two molecular partners (Spike and Furin) were found within 4.0 Å of each-other. In a salt-bridge thus formed, the positively charged nitrogen refers to side chain amino groups (-NH_3_ /=NH_2_ ) of lysines / arginines / doubly protonated histidines (His+) while the negatively charged oxygen refers to side chain carboxylic groups (-COO^-^) of glutamates / aspartates.

### 2.5.2. Analyzing salt-bridge dynamics

#### 2.5.2.1. Salt-bridge Persistence and Occurrence

Following standard analytical methods [49, 50], first, all unique salt-bridges occurring at least once in the MD simulation trajectories were identified and accumulated. The dynamic persistence (*pers*) of each unique (non-redundant) salt-bridge was then calculated as the ratio of the number of structural snapshots to which the salt-bridge was found to form with respect to the total number of snapshots sampled (at regular intervals) in the trajectory. As has been already mentioned (section **2.4**) an interval of 10 ps was chosen, leading to 30000 snapshots for the 300 ns trajectory (RR1_CoV-2_) and 10000 snapshots for the 100 ns trajectories (ZR1_CoV-2_: cross-validation, ZR1_CoV_: baseline). Likewise, for the static ensemble compiled of the top ranked 100 docked poses (RR1_CoV-2_ to RR100_CoV-2_), a static equivalent of persistence, namely, occurrence (*occ*) was procured for each salt-bridge occurring at least once in the ensemble using an equivalent ratio to that of the dynamic persistence (*pers*). Occurrence (*occ*) for each salt-bridge was defined as the ratio of docked poses to which the salt-bridge had occurred with respect to the total number of selected docked poses (=100) in the ensemble. Even a single occurrence of a salt-bridge in an ensemble was considered accountable in this analysis. Normalized frequency distributions of salt-bridge persistence (and occurrence) were plotted for the corresponding ensembles for further analyses.

#### 2.5.2.2. Average Contact Intensities of salt-bridges

To take care of the variable degrees of intensities (effectively, ionic strengths) of atomic contacts for salt-bridges, contact intensities (CI) were defined and computed for each salt-bridge as the actual number of inter-atomic contacts involved in a salt-bridge. In other words, CI is the number of ion-pairs to be found within 4Å between the two interacting side chains in a salt-bridge. Considering all unique combinations of possible salt-bridges (Arg ↔ Glu, Lys ↔Asp etc.), CI can vary from 1 to 4. Time series averages, defined as the average contact intensity (ACI) of these salt-bridges were then computed for each non-redundant salt-bridge from the MD simulation trajectories pertaining to each subject under test (RR1_CoV-2_, ZR1_CoV-2_, ZR1_CoV_). Together persistence (*pers*) and average contact intensity (ACI) can be considered as ‘ensemble descriptors of salt-bridges’. To account for their cumulative contribution in terms of salt-bridge strength and sustenance, a weighted persistence term (*wpers(i)*) was further defined for each i^th^ non-redundant salt-bridge in an ensemble, as the direct product of *pers(i)* and ACI(i).

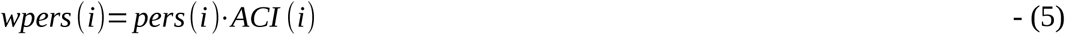

By definition, *wpers* would have a theoretical range of [0,4].

Furthermore, in order to draw a direct comparison between cumulative contact intensities (CCI) of the ionic bond networks formed across the different Spike–Furin interfaces in SARS-CoV (ZR1_CoV_) and SARS-CoV-2 (RR1_CoV-2_, ZR1_CoV-2_) the following sum-over measure was defined, designed and implemented.

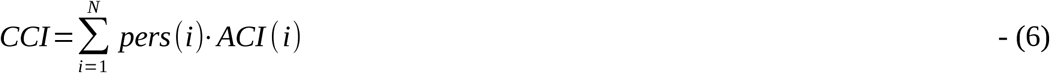

- where, *pers(i)* and *ACI(i)* were defined as before (in Eqn. 5) and N is the total number of non-redundant salt-bridges found at least once in a dynamic ensemble. In the current context, CCI can be considered as a measure of structural degeneracy [88] that is made to function as a global network descriptor raising a limiting threshold at the coronavirus Spike–Furin interfaces, allowing, absorbing and accommodating different alternative ionic bond network architectures as long as they are befitting to the task of catalyzing the Spike–S1/S2 cleavage.

### 2.6. Calculation of structure-based equilibrium thermodynamic parameters (ΔH, ΔS, ΔG) for the Spike–Furin binding

As a mean to probe a highly likely event of ‘enthalpy – entropy compensation’ associated implicitly with the the Spike–Furin interaction, structure-based equilibrium thermodynamic parameters (ΔH, ΔS, ΔG) were calculated for the selected representative structure (RR1_CoV-2_) along its entire MD simulation trajectory (300 ns) using the standalone (C++ with boost library) version (v.4) of FoldX (http://foldxsuite.crg.eu/) [89, 90]. FoldX has its energy terms carefully parameterized by actual experimental data from protein engineering studies [89] – which together with its high computational speed are definite edges over the traditional MM(PB/GB)SA approaches [91]. It is for these reasons, FoldX is slowly but surely taking over traditional approaches in structure-based thermodynamic calculations, particularly in the domain of protein engineering and stability analysis [92, 93]. FoldX is built on a ‘fragment-based strategy’ that exploits the power of fragment libraries [94] in the same direction to that of the most compelling ‘fragment assembly simulated annealing’ approach in protein structure prediction attributed to David Baker and Rosetta [64]. Along with net free-energy changes (ΔGG_binding/folding_) the advanced empirical force field of FoldX also returns a plethora of different favorable or disfavored transition enthalpic as well as entropic energy terms for proteins (folding) and PPI complexes (binding) directly from their high-resolution 3D coordinates (using full atomic description). To address the plausible ‘enthalpy – entropy compensation’ in the current context, as enthalpic terms we included the favorable van der Waals (ΔH_vdwF_) and electrostatic (ΔH_electro_, ΔH_elec-kn_) contributions to free energy, as well as the disfavored van der Waals clashes (ΔH_vdw-clash_, ΔH_vdw-clash-backbone_); while, to account for the entropic costs, we included the entropic energies for backbone (TΔS_mc_) and side chain (TΔS_sc_) conformational changes. The choice of the terms was guided by well-just reviews and discerning followup studies on FoldX [92, 95]. The enthalpic terms were further summed up according to the nature of forces giving rise to each.

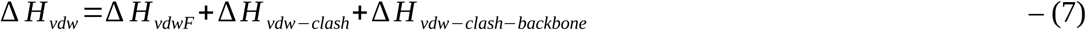

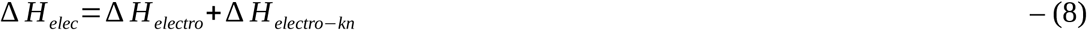

To that end, structural snapshots were sampled at 10 ps interval from the 300 ns MD simulation trajectory (RR1_CoV-2_) resulting in 30000 time-stamps (or, structural snapshots). Then, for each snapshot, FoldX was run using the command *AnalyseComplex* with the *complexWithDNA* parameter set to ‘false’ and the relevant enthalpic (ΔH_vdw_, ΔH_elec_), entropic (TΔS_mc_, TΔS_sc_) and free energy terms (ΔGG_binding_), as detailed above, were computed for each run, indexed appropriately and stored. Time averages (denoted by angular braces ‘<>’ throughout the paper) were computed for each individual term along with its standard deviations (SD). For a second analyses, focusing purely on the ‘entropy arrest’ [96–98] presumably implicit to the Spike–Furin binding, conformational entropies for backbone (subscripted as ‘mc’) and side chains (subscripted as ‘sc’) were recorded (for each time-stamp) independently for the FoldX–separated (unbound) receptor (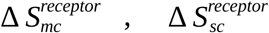) and ligand (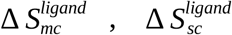), as well as in their bound forms (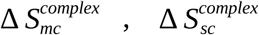 Lastly, from the individual time-series averages of the ΔG_binding_ values obtained for the Spike– Furin binding in SARS-CoV and SARS-CoV-2, ΔΔG_binding_ was defined as follows taking care of the cumulative effect of mutational changes at their FLCS_Spike_.

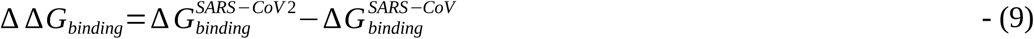

### 2.7. Ramachandran Plot (RP) - derived parameters to probe state-transitions (e.g., disorder-to-order)

The Ramachandran Plot (RP) [51] can effectively be used to probe transitions between disordered and relatively ordered protein structural elements. To achieve this, first, dynamic conformational ensembles need to be assembled, representative of two protein states, say, an unbound (highly disordered) and a bound (relatively ordered) state. Since, RP is essentially based on local steric clashes, the analysis can further be locally restricted to a ‘contiguous region of interest’ (or, a concerned structural patch) wherein residues are supposed to undergo a two-state transition (say, *disorder-to-order*). In the current study, this ‘contiguous region of interest’ is the FLCS_Spike_ loop and the two protein states are the unbound (disordered) and Furin-bound (presumably more ordered) states of the SARS-CoV-2 Spike. The Ramachandran angles (*Φ, ψ*) can then be computed for residues comprising the concerned structural patch and plotted in an RP for each atomic model / frame in an ensemble state. The individual RPs can then be overlaid for ‘disordered’ or ‘relatively ordered’ states. In order to identify whether a connected structural patch (especially, relevant for protein loops) supports a finite set of (restrained) structural conformations, the 2-dimensional euclidean distance in the *Φ*–*ψ* vector space of RP were computed for each internal residue comprising the concerned structural patch. The distance between Ramachandran angles of i^th^ and j^th^ residues in *Φ*–*ψ* space can be interpreated in terms of the extent of local conformational mismatches between i^th^ and j^th^ residues and may be formally represented as follows:

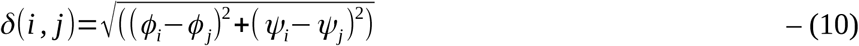

The maximum value of δ(i,j) represents the maximum spread of (*Φ, ψ*) within the concerned structural patch. It naturally follows that the concerned patch is structurally relatively more ordered when this spread is comparatively less and vice-versa. For some statistical reference, say <δ>_9d_ be the 9^th^ decile of δ(i,j). This indicates that 90% of the residues are separated by some distance less than <δ>_9d_ in the {*Φ, ψ*} space. Consequently, higher structural order is reflected through lesser values of <δ>_9d_ which increases with the increasing disorder in the structure. We took three statistics from the distribution of these distances δ(i,j) obtained for each state: (i) the median (50 percentile, <δ>_median_), (ii) the 3^rd^ quartile (75 percentile, <δ>_3q_) and (iii) the 9^th^ decile (90 percentile, <δ>_9d_) which are adequate to collectively render a comparison across states. Use of such a combined statistics instead of the maximum value of δ(i,j) also implicitly avoid possible outlier effects.

It is also necessarily important to estimate the local coherence of structural conformation within the concerned structural patch in general. Such a measure (metric) could be informative in terms of the local tendencies within (say) a protein loop to attain certain restrained local structural conformations. To estimate local structural coherence within the concerned structural patch, the average euclidean distance (along with the standard deviation) of (*Φ, ψ*) points between consecutive residues comprising the concerned structural patch was designed and computed in the following way:

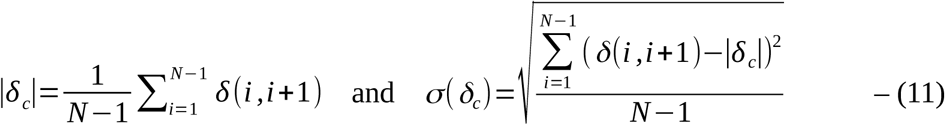

where |δc| and σ(δ_c_) are defined for each atomic model / frame falling within an ensemble (state) and the ‘contiguous region of interest’ (or concerned structural patch) is N-residues long. Relative lower values of |δ_c_| represents higher local structural coherence and σ(δ_c_) presents a measure of dispersion in local structural coherence within a protein loop. Hence, these ordered parameters may be computed to collectively render a comparison of local structural coherence across states.

### 2.8. Quantifying a change between two N-binned frequency distributions and assessing its statistical significance in terms of χ^2^

A χ^2^ test (wherever applicable) was conducted to discriminate between two frequency distributions (say, that of an unbound and a bound state of a protein region spanning different contoured regions the RP) with the χ^2^-statistic being computed (for an N-bin model; df^4^=N-1) by the following equation.

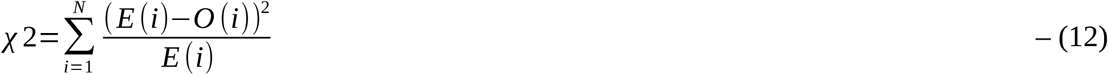

- where *E(i)* represents the frequency ‘under the null hypothesis’ expected for the i^th^ bin, while, O(i) denotes the actually observed frequency for that same (i^th^) bin.

## 3. Results and Discussion

### 3.1. Structural insight into the Furin cleavage mechanisms

From a structural, biochemical as well as from a biophysical perspective, it is crucial to unravel the reaction mechanisms and the involved enzyme kinetics of the Furin cleavage demanding quantum chemical and/or QM/MM^5^ studies. To that end, a preceding step would be to explore the nature of binding involved in the Spike–Furin interactions via the disordered FLCS_Spike_ - loops in SARS-CoV and SARS-CoV-2, compare them and contrast. To address this, here we adapted a combined approach of ensemble molecular docking and dynamic simulations followed by conformational analyses. As briefed in the **Introduction**, the coronavirus Spike structures are devoid of experimental atomic coordinates for the FLCS_Spike_ patch which is further revealed to be a “surface exposed disordered loop” [21] for the pre-fusion structure of SARS-CoV-2 Spike. There are as many as 44 experimentally solved (exclusively cryo-EM) structures (**Table S1, Supplementary Materials**) currently to be found in the PDB (see section. **2.1.2**, **Materials and Methods**) for the pre-fusion form CoV/CoV-2 Spike, culled at a resolution of not worse than 3 Å. These coronavirus Spike structures are solved to serve different specific research objectives at various pH [99] and other varying physico-chemical conditions [17,58,60,61], therefore, often requiring stabilizing (engineered) mutations at the FLCS_Spike_ patches [24, 38]. It is almost intriguing that even with stabilizing modulations, there’s not a single cryo-EM structure that has any experimental (primary) data for its FLCS_Spike_ patch. As a result, atomic coordinates of the *_681_PRRAR_685_* motif in the SARS-CoV-2 Spike (and, equivalent homologous sequence motifs from other coronavirus Spike) along with short flanking regions at both ends (adding up to a stretch of 10-12 amino acids) are missing experimentally [39] and hence require computational modeling (**Figure S2**, **Supplementary Materials**). To have such disordered loops appears quite characteristic of the SARS-CoV-2 Spike trimer which contains a total of four missing stretches of roughly similar lengths at strategic positions, adding up to 42 amino acids in PDB ID: 6XR8. This is perhaps reasonable given the extended internal packing involved in the trimerization of the monomeric S units. The highly localized positive charge cloud concentrated over the arginine-rich _681_PRRAR_685_ region of the loop further boosts the said probability as this would instigate electrostatic self-repulsion of the loop adding to its conformational instability. The presence of Proline (P681), the well-known helix breaker, within the SARS-CoV-2 FLCS_Spike_, plausibly adding to the loop-disorder, has further been suggested to improve the protease active site accessibility for Furin as well as for other proteases [100].

### 3.2. ‘More the arginines, more the disorder’ in the FLCS_Spike_ activation loops

In order to have a general idea as to how the presence of polybasic sequences (arginines) influence the evolutionarily manifested disorder in the FLCS_Spike_, we started the proceedings with an evolutionary analysis of the loop-disorder on compiled coronavirus Spike sequences. There are several AI^6^-trained sequence based disorder predictors [101–103] that return residue-wise disorder probability scores, which are trained primarily on evolutionary sequence data (e.g., mutational co-variance matrices). These sequence-based disorder predictors have their known limits in accuracy [104, 105], for not explicitly accounting for the actual three-dimensional structural dynamics of the protein(s) / peptide(s), but, can serve as a good first test of the comparative FLCS_Spike_ loop-disorder among its close evolutionary homologs. A representative set of Spike structures (CoV/CoV-2) were culled (resolution ≤ 3 Å), accumulated (see **subsection 2.1**, **Materials and Methods**), and their UNIPROT sequences (in FASTA format) derived from proteomics data [63], were extracted from corresponding entries in the Protein Data Bank [39]. The full-length Spike sequences were aligned using MUSCLE [106] and those containing gap(s) at their aligned position(s) homologous to the _681_PRRAR_685_ pentapeptide motif (FLCS_Spike_, SARS-CoV-2) were removed. The final set consisted of all unique and non-redundant pentapeptide sequence motifs (PSGAG, PGSAS, PASVG, PSRAG, PSRAS, PRRAA, PRARR) to be found within the FLCS_Spike_ spanning the entire plethora of coronavirus Spikes. The full-length sequences were then run in the PrDos web-server [102] – which combines local sequence information and homology templates using iterative (psi) BLAST. Since the loop-disorder is highly contextual to its neighboring / flanking sequences and to that of the ‘highly conserved’ trimeric Spike structures (C^α^-RMSD: 2.3 Å), the default setting of ‘template-based’ prediction (with the PrDos-FPR^‡7^ set to 5%) was retained as ‘turned on’. For all representative full-length Spike sequences, the disorder probabilities of the FLCS_Spike_ regions (i.e., the patches originally missing in the corresponding experimental structures) were unanimously found to increase sharply in the N→C direction around the pentapeptide motifs trending to local maximums (**Figure 1**). The regions, consequently mapped to the ascending halves of the corresponding curve-humps (**Figure 1.A**) – which should effectively mean ‘growing disorders’ associated with the FLCS_Spike_. Interestingly, the mean disorder probabilities have an unmistakable increasing trend (0.45 for PSGAG →→ 0.55 for PRARR) with the successive gradual incorporation of arginines in the pentapeptide sequence motif.

**Figure 1.**
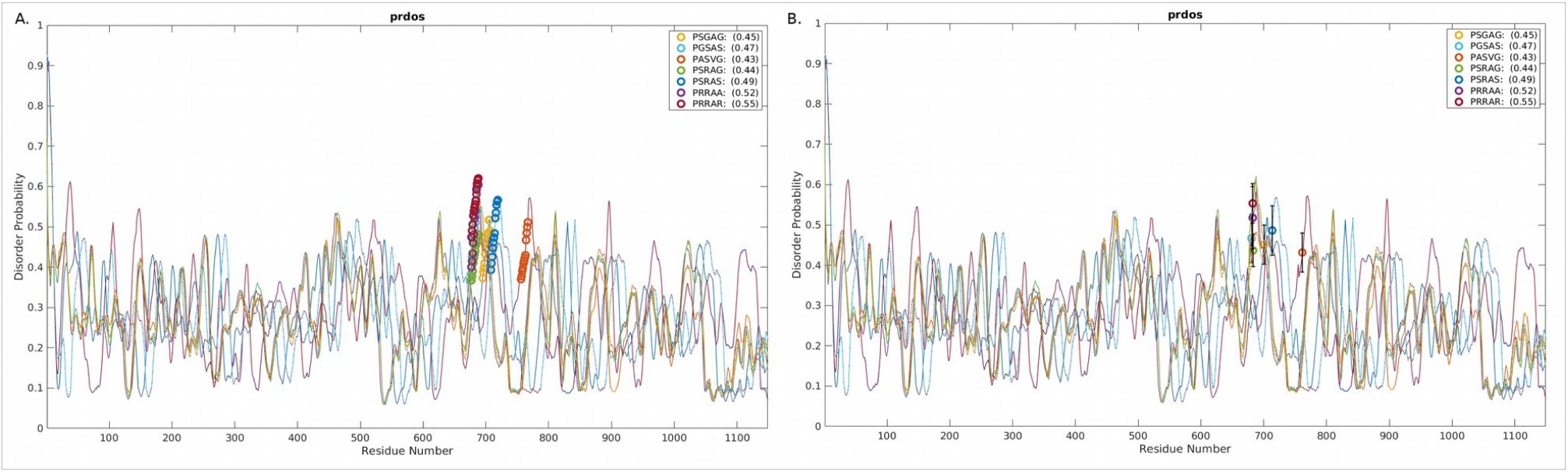
PrDos disorder probability scores plotted for coronavirus Spike sequences. The sequences cover the entire non-redundant sequence space up to the evolution of SARS-CoV-2 for the pentapeptide sequence motifs (e.g., *_681_PRRAR_685_* in SARS-CoV-2) embedded in FLCS_Spike_. Panel (A) portrays the residue-wise disorder probabilities (with the FLCS_Spike_–residues highlighted with deferentially colored thick circles) while panel (B) plots their mean values centered on the middle-most residue in the corresponding FLCS_Spike_. The error bars represent the corresponding standard deviations.

### 3.3. Filling up the voids in the Spike structures: the FLCS_Spike_ disordered ensembles

The disorder, intrinsic to the FLCS_Spike_ along with the most likely event of ‘*disorder-to-order* transition’ upon binding to Furin (see **Introduction**) makes the Spike–Furin iteration an interesting mechanistic chapter both in the context of coronavirus evolution and also in the general framework of proteolytic cleavages [41–43]. Particularly intriguing (and, counter-intuitive almost) is the fact that the bait the SARS-CoV-2 Spike uses to attract their specific and dedicated host-proteases (e.g., Furin for SARS-CoV-2 Spike) are themselves structurally highly nonspecific or conformationally varied by virtue of their intrinsic disorder. Again, in the bound form, the concerned disordered loop (FLCS_Spike_, SARS-CoV-2) serves as the bait to recruit Furin subsequent to which it needs to be jammed into a much restricted set of conformations for its efficient (proteolytic) cleavage [32]. The collective dynamics of the Spike–Furin interaction (SARS-CoV-2) would thus naturally lead to a conformational selection of the disordered FLCS_Spike_ in its Furin-bound state. To sustain such an energetically costly ‘conformational selection’, an energy-source is hence required to compensate for such a high entropy-loss of the disordered FLCS_Spike_ ensemble. Given the physico-chemical nature of long- and short-range forces sustaining the native bio-molecular environment, such compensating energies would naturally be enthalpic in nature. Thus, an enthalpy – entropy compensation seems necessary. So, the obvious question that follows next would be as to what might be the source of this enthalpic energy to compensate for such a high entropy-loss?

To address this, first and foremost, the missing disordered stretches in the representative SARS-CoV-2 Spike pre-fusion structure (PDB ID: 6XR8) were ensemble-modeled (see, section **2.2, Materials and Methods**) using its full-length amino acid sequence, derived from proteomic data. Considering that the SARS-CoV-2 FLCS_Spike_ (6XR8) is only a short stretch of 12 amino acid residues (_677_QTNS**PRRAR**SVA_689_), 500 uniquely varied loop-conformations were sampled for the missing patch, eventually leading to that many ‘all-atom’ trimeric SARS-CoV-2 Spike atomic models. Such an ensemble can essentially and adequately represent the conformational variations in the disordered SARS-CoV-2 FLCS_Spike_ (missing) patch. During the entire course of this modeling, the atoms that were already experimentally solved were retained as they were. The average C^α^-RMS deviations, upon aligning the models pairwise in PyMol (https://pymol.org/2/<u>), were found to be appreciably higher (4.71 Å; SD: 0.82) even for the short (101 atoms) modeled FLCS_Spike_ – loop (resi. 677-688) compared to the ‘all-atom’ Spike structures (2.93 Å; SD: 0.54 for 26940 aligned atoms). This further confirmed the disorder, intrinsic to the FLCS_Spike_ in SARS-CoV-2. To serve as a baseline, a similar approach was adapted to model the homologous missing patch (_664_SLLRSTS_670_) in SARS-CoV Spike (see,</u> section **2.2<u>, Materials and Methods</u>**<u>) that was originally missing in the representative experimental structure (PDB ID: 7AKJ), albeit with a lower degree of conformational sampling proportional to a lower degree of disorder (than that of SARS-CoV-2) in its FLCS_Spike_ (see,</u> section **2.2<u>, Materials and Methods</u>**).

### 3.4. Docking Furin onto Spike: using the pentapeptide activation loop to filter and accumulate correctly docked poses

Once the (unbound) disordered FLCS_Spike_ ensemble (SARS-CoV-2) was made ready (section **3.3**), a series of blind docking experiments was conducted in ClusPro 2.0 [65] (see **2.3.1, Materials and Methods**), wherein Furin (ligand) was docked *ab-initio* onto each of the 500 Spike ‘all-atom’ atomic models (receptors) without imposing any additional ‘active-site / contact residue’ constraints. This resulted in an initial pool of 53215 Spike–Furin unsorted docked poses (**Figure S3, Supplementary Materials**) with the ligand docked at widely varying sites spread all over the trimeric Spike receptor. The extra-large sample space of returned docked poses (of the order of fifty thousands) ensured necessary and sufficient coverage of the ligand – receptor orientational space in docking while the detailed iterative blind docking protocol (Cluspro 2.0) maintained the fine-grain qualities of the docked poses. All docked poses for each ensemble- docked set (consisting of 500 receptor templates unique for their FLCS_Spike_) were then accumulated. Since the ‘desired interface’ would necessarily involve the FLCS_Spike_, the pentapeptide motif (_681_PRRAR_685_) was used as ‘contact residue filters’ to discard obviously and/or trivially incorrect docked poses. To that end, buried surface areas (BSA) (see section **2.3.2.1**, **Materials and Methods**) were computed for all residues in each docked pose and the interfacial residues (BSA≠0, see section **2.8**) were identified. In addition, the BSA values for residues pertaining to the _681_PRRAR_685_ motif (BSA_PRRAR_) were summed up and stored for each docked pose. If the interfacial residues, so identified, happen to contain any fraction of the pentapeptide motif (i.e., BSA_PRRAR_>0), the docked pose was considered plausible and worthy to be carried for the next round (i.e., filtered in). All three chains of the Spike – trimer (harboring this FLCS_Spike_) were treated equally likely to serve as the docking site. This simple technique ensured that all filtered docked poses (7184 of them) have the desired Spike–Furin interface harboring the FLCS_Spike_ [48]. It was interesting to note that in the top-ranked docked poses (ranked by Cluspro’s internal score) for almost all ensemble-docked batches (in 497 out of 500 cases) Furin had an unmistakable preference to dock at the FLCS_Spike_ (i.e., BSA_PRRAR_>0) even with this unbiased *ab-initio* blind docking protocol.

After the initial filtering of ‘plausible’ docked poses (those that harbors the FLCS_Spike_ at the Furin – Spike interface), Sc^FLCS^ (see section **2.3.2.2, Materials and Methods**) was computed for the BSA – filtered interfaces alone. To render a discerning scoring function appropriately combining both surface fit and overlap at the interface, BSA_PRRAR_ was further normalized to nBSA_PRRAR_ (see section **2.3.2.1, Materials and Methods**). A second subsequent filtering was performed on the filtered set wherein nominal relaxed cutoffs (slightly more restraint than merely non-zero) on nBSA_PRRAR_ and Sc^FLCS^ were jointly implemented (nBSA_PRRAR_>0.05, Sc^FLCS^>0.2) based on their general trends [71, 77]. In effect, each docked pose had to attain minimum thresholds in terms of both the goodness of fit of the interacting surfaces and their extent of conjointness. The second round of filtering thus resulted in more realistic poses and further reduced their number from 7184 to 5735. These poses were then scored and re-ranked by an appropriately weighted scoring function (S_dock_) combining nBSA and Sc (see section **2.3.2.3, Materials and Methods**). The re-ranking ensured careful and combined monitoring of both the geometric fit of the interacting surface patches at the interface and also the extent of that fit. The average {<nBSA_PRRAR_>, <Sc^FLCS^>} values for the top 10 (re-)ranked docked poses (**Table S2, Supplementary Materials**) were suggestive of their elevated surface fit (<Sc^FLCS^>: 0.70, SD^8^: 0.04) and decent surface overlap (<nBSA_PRRAR_>: 0.211, SD: 0.034) at the interface. The same averages for the top 100 (re-)ranked docked poses were 0.66 (SD: 0.06), 0.129 (SD: 0.053) for Sc^FLCS^, nBSA_PRRAR_ (**Table S2, Supplementary Materials**) – which fall very well within the range of values obtained for the same or equivalent parameters surveyed at native protein interiors [72–74] as well as at native/near-native protein interfaces [71, 77]. Top ranked docked poses were all visually surveyed and scrutinized in PyMol. No noticeable inconsistencies were observed. The top ranked docked pose (RR1_CoV-2_) with a S_dock_ score of 0.972 was also ranked first in terms of surface overlap (BSA_PRRAR_: 546.128 Å^2^; nBSA_PRRAR_: 0.247) and third in terms of shape complementarity (Sc^FLCS^: 0.77) to be attained at the Spike–Furin interface. RR1_CoV-2_ was then taken to energy minimization cycles followed by a (sufficiently long) explicit water, all atom molecular dynamic simulation (see, section **2.4, Materials and Methods**) to account for the structural dynamics of the Spike–Furin interaction in SARS-CoV-2.

In contrast, a guided docking approach using ‘Zdock + IRaPPA re-ranking’ (see sections **2.3.1.2**, **2.3.1.3**, **Materials and Methods**) was directly adapted in parallel as a mean to cross-validate the results obtained from RR1_CoV-2_ as well as for the baseline subject in SARS-CoV leading to two more top-ranked models, ZR1_CoV-2_, ZR1_CoV_ respectively – both of which were then undertaken long (100 ns) MD simulation runs, subsequent to equivalent rounds of energy minimization.

### 3.5. Plausible ‘disorder-to-order’ transition triggered by salt-bridge dynamics at the Spike– Furin interface: the ‘Salt-bridge hypothesis’

One common intuition to jam the high-entropy FLCS_Spike_ disordered loop (SARS-CoV-2) into the Furin bound enzyme-substrate complex would be to stabilize the highly localized positive charge cloud electrostatically. It is rather well known that electrostatic interactions play an important role in the dynamic sustenance and transitions associated with protein disorder [107–112]. To that end, the suggested electrostatic stabilization of the FLCS_Spike_ would greatly be benefited by the structural proximity of oppositely charged (i.e., anionic) amino acids (coming from Furin) by triggering the formation of Spike–Furin interfacial salt-bridges. The plausibility of the ‘salt-bridge hypothesis’ is further enhanced by the presence of a surface groove around the Furin [32] catalytic triad (D153, H194, S368) that appears to be befitting to FLCS_Spike_ both in terms of shape and electrostatics (**Figure 2**). This groove serves as a potentially attractive docking site evolutionarily for FLCS_Spike_ patches. A detailed independent survey of the two-domain Furin structure (PDB ID: 1P8J, chain A, see section **2.1.1**, **Materials and Methods**) further reveals that it has an abundance of anionic residues (30 aspartates and 25 glutamates) which are spread rather homogeneously all over its catalytic and P domains (see **Introduction**). While the majority (58%) of these residues are either partially or completely exposed to the solvent (*bur*≥0.05) (see section **2.3.2.1**, **Materials and Methods**), those that are part of the catalytic domain (**Figure 2**) are of interest to the given context. Note that even partially exposed anionic residues with lesser exposure (say, 0.05<*bur≤*0.15) in vicinity of the triad can, in principle, seldom flip into a more extended and exposed conformation during the course of the Spike–Furin binding (till the Spike–S1/S2 cleavage). Hence, these may also potentially contribute to stabilize the proposed dynamically interchangeable ionic bond networks at the Spike–Furin interface (SARS-CoV-2). To have a closer look into these ‘residues of interest’, first, a subset of anionic residues in Furin was assembled, where at least one heavy side chain atom from each residue is located (in its crystalline equilibrium state) within a sphere of radius 20 Å centered at the side chain centroid of the Furin catalytic triad (D153, H194, S368). This subset contained 25 residues, most of which (18 out of 25) are ‘partially exposed’ or ‘exposed’ (**Table S3**, **Supplementary Materials**) and hence should be amenable and approachable for formation of interfacial salt-bridges. Again, some of these partially or completely exposed anionic residues (viz., D191, D228, E236, E257, D258, D259, D355, D306; see **Table S3**, **Supplementary Materials**) are rather proximal to the catalytic triad (within 15 Å from its side chain centroid). Together, these constitute a non-rigid set of anionic Furin residues that potentially hold the FLCS_Spike_ at the catalytic triad, with the extent of stability required to encompass the cleavage, by virtue of dynamically engaging themselves into ionic bond interactions with the FLCS_Spike_–arginines coming from _681_PRRAR_685_. The positioning of these anionic side chains also appears to be strategic (**Figure 2**) such that cumulatively the salt-bridges almost never get dissolved during the entire course of the Spike–Furin binding. Such dynamic networks of potentially interchangeable ionic bonds could also be a prime source for the enthalpy required to compensate for the presumably high entropic loss of the FLCS_Spike_ upon binding to Furin. The process would also necessarily accompany a concomitant transition of the FLCS_Spike_ from a disordered (unbound) to a relatively ordered (Furin bound) state.

**Figure 2.**
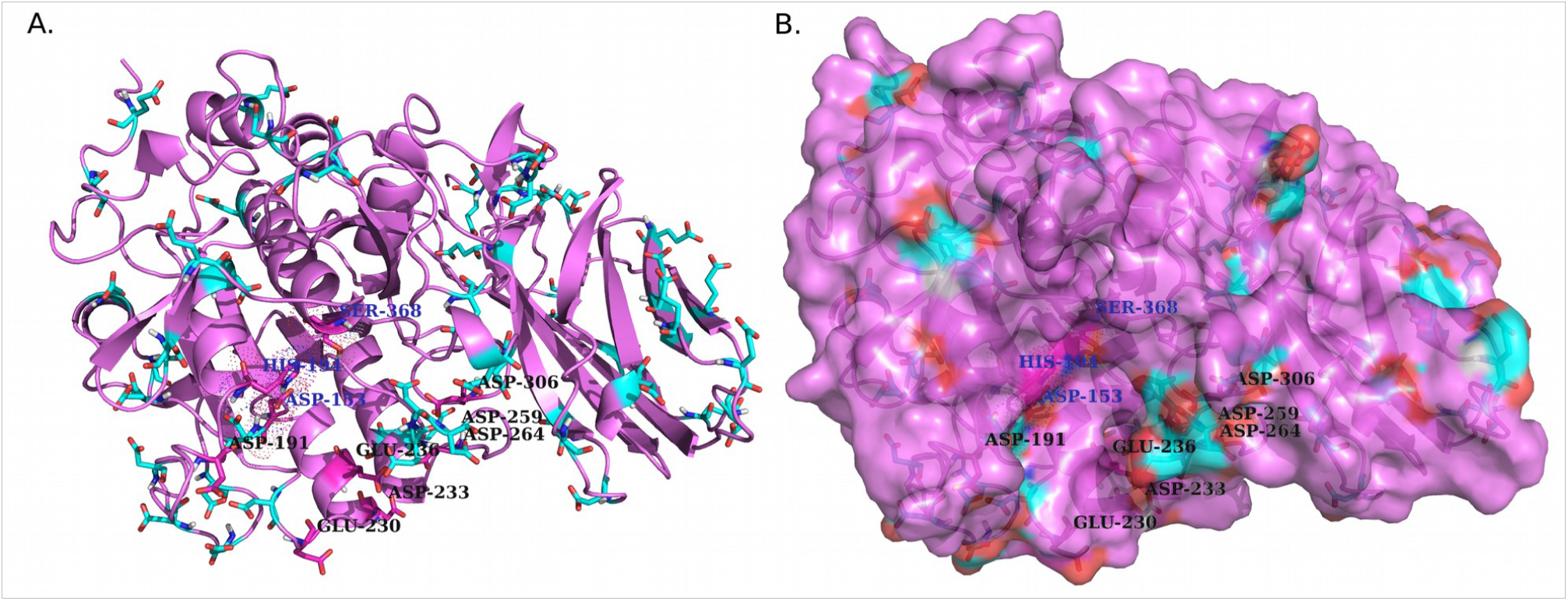
The proposed dockable surface groove for FLCS_Spike_ in Furin, proximal to its catalytic triad. The catalytic triad (D153, H194, S368) is highlighted dots and labeled in PyMol color: ‘density’ (panel **A**) over and above the cartoon display. All exposed and partially exposed anionic residues are shown in ‘balls and sticks’. Among these, the ones that are in close vicinity to the dockable groove surface (panel **B**) are further labeled with residue identities in both panels. These residues are amenable to form interfacial salt-bridges with the FLCS_Spike_–arginines (in _681_PRRAR_685_) and collectively add to the potential of forming dynamically interchangeable ionic bond networks at the Spike–Furin interface.

### 3.6. Validations and cross-validations of the ‘salt-bridge hypothesis’

#### 3.6.1. In RR1_CoV-2_

In order to test the validity of the ‘salt-bridge hypothesis’, an all-atom explicit water MD simulation (of 300 ns) was run (see section **2.4**, **Materials and Methods**) for the top ranked Spike–Furin docked pose (RR1_CoV-2_), and, the trajectories were tested for the presence of salt-bridges (see section **2.5.1**, **Materials and Methods**). Prior to the production run, the structural template (RR1_CoV-2_) was undertaken rigorous rounds of energy-minimization with temperature equilibration (see section **2.4**, **Materials and Methods**). Structural snapshots were sampled at 10 ps (regular) intervals leading to 30000 snapshots (or time-stamps) accounting for the 300 ns long MD simulation trajectory and salt-bridges were extracted from each snapshot from its interfacial contact map (see section **2.5.1**, **Materials and Methods**). To set a reference, ionic bond networks were also surveyed in the pool of 100 top ranked static docked poses (**Table S2**, **Supplementary Materials**). In both ensembles, dynamic and static, the identification of salt-bridges were followed by their individual survey and also a statistical analyses of dynamic or static ‘ensemble descriptors of salt-bridges’ (e.g., persistence/occurrence and average contact intensities; see section **2.5.2**, **Materials and Methods**). Together these parameters can effectively be used to decipher and interpret the complex nature of salt-bridge dynamics associated with protein disorder transitions [49, 50]. One of the trademark features of salt-bridge dynamics associated with disorder transitions is a trade-off or an optimal balance between transience and persistence of salt-bridges which serves to sustain a disordered state while a shift of this balance leads to disorder transitions [49, 50].

To that end, explicit lists of ionic bond interactions independently for the dynamic (**Table 1**), static ensembles (**Table S4, Supplementary Materials**) were sorted based on their persistence (dynamic), occurrence (static) (see section **2.5.2.2., Materials and Methods**). Given that the static ensemble consisted of near-native as well as less near-native poses (**Figure S4**, **Supplementary Materials**), it was interesting to find that all (or almost all) Spike–Furin interfacial salt-bridges that had occurred (at least once) in the static ensemble were also found (at least ephemerally) in the dynamic ensemble. The persistence/occurrence profiles collectively reveal that the Spike–Furin interaction has a preferential set of ionic bonds in terms of forming dynamically interchangeable salt-bridges – which may vary in their occurrence among the plausible docked poses. Since, RR1_CoV-2_ presents possibly the most preferred conformation of the FLCS_Spike_ in its Furin bound state (in SARS-CoV-2), the dynamically sustained high persistence arginine – salt-bridges (in _681_PRRAR_685_) found in RR1_CoV-2_ are the most plausible, frequently forming Spike–Furin interfacial salt-bridges (**Figure S4**, **Supplementary Materials**) among alternatives.

**Table 1.**
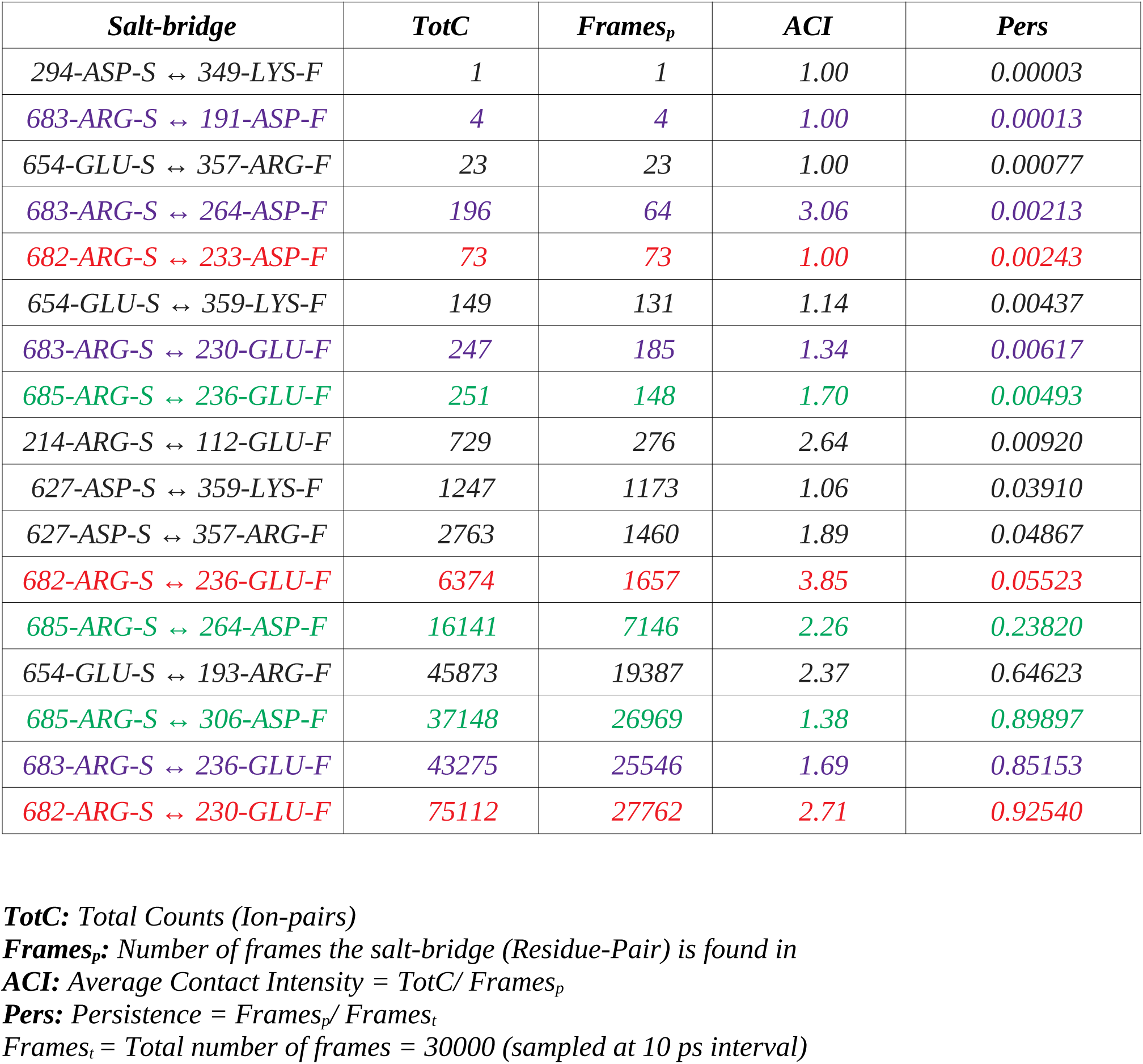
Persistence and average contact intensities of all unique salt-bridges at the SARS-CoV2 Spike–Furin interface for RR1_CoV2_ (the top re-ranked docked pose) along its 300 ns MD simulation trajectory. ‘-S’ & ‘-F’ in the salt-bridge descriptor strings refer to the receptor and the ligand chains respectively. Rows corresponding to the Arginine-salt-bridges falling within the pentapeptide _681_PRRAR_685_ motif (activation loop of FLCS_Spike_) is highlighted in three different font colors for R682, R683, R685.

| *Salt-bridge* | *TotC* | *Frames<sub>p</sub>* | *ACI* | *Pers* |
| --- | --- | --- | --- | --- |
| 294-ASP-S ↔ 349-LYS-F | 1 | 1 | 1.00 | 0.00003 |
| 683-ARG-S ↔ 191-ASP-F | 4 | 4 | 1.00 | 0.00013 |
| 654-GLU-S ↔ 357-ARG-F | 23 | 23 | 1.00 | 0.00077 |
| 683-ARG-S ↔ 264-ASP-F | 196 | 64 | 3.06 | 0.00213 |
| 682-ARG-S ↔ 233-ASP-F | 73 | 73 | 1.00 | 0.00243 |
| 654-GLU-S ↔ 359-LYS-F | 149 | 131 | 1.14 | 0.00437 |
| 683-ARG-S ↔ 230-GLU-F | 247 | 185 | 1.34 | 0.00617 |
| 685-ARG-S ↔ 236-GLU-F | 251 | 148 | 1.70 | 0.00493 |
| 214-ARG-S ↔ 112-GLU-F | 729 | 276 | 2.64 | 0.00920 |
| 627-ASP-S ↔ 359-LYS-F | 1247 | 1173 | 1.06 | 0.03910 |
| 627-ASP-S ↔ 357-ARG-F | 2763 | 1460 | 1.89 | 0.04867 |
| 682-ARG-S ↔ 236-GLU-F | 6374 | 1657 | 3.85 | 0.05523 |
| 685-ARG-S ↔ 264-ASP-F | 16141 | 7146 | 2.26 | 0.23820 |
| 654-GLU-S ↔ 193-ARG-F | 45873 | 19387 | 2.37 | 0.64623 |
| 685-ARG-S ↔ 306-ASP-F | 37148 | 26969 | 1.38 | 0.89897 |
| 683-ARG-S ↔ 236-GLU-F | 43275 | 25546 | 1.69 | 0.85153 |
| 682-ARG-S ↔ 230-GLU-F | 75112 | 27762 | 2.71 | 0.92540 |

**TotC:** Total Counts (Ion-pairs)
**Frames<sub>p</sub>:** Number of frames the salt-bridge (Residue-Pair) is found in
**ACI:** Average Contact Intensity = TotC/ Frames<sub>p</sub>
**Pers:** Persistence = Frames<sub>p</sub>/ Frames<sub>t</sub>
Frames<sub>t</sub> = Total number of frames = 30000 (sampled at 10 ps interval)

The first noticeable observation in the dynamic ensemble (pertaining to RR1_CoV-2_) was that the three arginines (R682, R683, R685) in the _681_PRRAR_685_ almost always remain engaged in dynamically interchangeable ionic bonds formed with anionic side chains coming from the host Furin. These amenable anionic side chains (D191, E230, D233, E236, D259, D264, D306, see **Table S4, Supplementary Materials**) are indeed physically proximal to the catalytic triad (**Figure 3.A**), nicely fitting in the FLCS_Spike_ into the proposed dockable surface groove. While most (if not all) of them fall into the non-rigid ‘expected’ set (**Table S3, Supplementary Materials**), they should collectively present some correlated movements with FLCS_Spike_ in order to remain conformationally viable for dynamically stable ionic bond networks. Further, as reflected from the distribution of salt-bridge persistence (see **Table 1**) the three arginines in _681_PRRAR_685_ were often involved in multiple and interchangeable ionic bonds. In other words, the same arginine (target) in _681_PRRAR_685_ can simultaneously be in contact (at a given instance) with more than one negatively charged residues coming from the host Furin (neighbor). This will lead to the formation of non-disjoint sets of target-neighbor ionic bond pairs (i.e., salt-bridges) for the same target. In other words, sets of salt-bridges having distinct anionic partners for a given target arginine (_681_PRRAR_685_) would thus be non-disjoint. This implies that the persistence values of the arginine – salt-bridges (in _681_PRRAR_685_) may add up to more than unity (see **Table 1**). In fact, dynamic interchangeability of counter-ions are common characteristic features of salt-bridges formed at disordered protein regions and/or [49, 50] disorder–globular interfaces [86, 110] wherein the formed ion pairs keep changing their counter-ionic partners [49]. Such collective dynamics results in salt-bridges of varying persistence and multiplicity across the trajectory, including persistent as well as transient salt-bridges [49,50,109,111]. Transient and/or unfavorable salt-bridges have been revealed to be functionally optimized in proteins [49, 112] and are often found on enzyme surfaces [112] as well as on strategic locations sparsed around extended disordered protein regions [49]. For the later case, one of the evolved key mechanisms towards achieving this functional optimization is to make their charged side chains often amenable to proximally approached ordered protein interactors. The transient nature facilitates the exchange of their counter-ionic partners, thereby triggering the switch from intra- to inter- molecular (interfacial) salt-bridges. The multiplicity (or promiscuity [49]) in the choice of the counter-ionic partner in a salt-bridge can further be chemically and electrochemically rationalized from the bifurcated nature of side chain groups with degenerate charge centers in four out of the five charged amino acids (guanidium^+^: Arg; carboxylate^-^: Asp, Glu; imidazol^+^: His+) in proteins. However, in spite of this observed multiplicity, each of the three arginines in _681_PRRAR_685_ seemed to have their own preference for particular anionic partners: E230 for R682 (*pers*: 0.93), E236 for R683 (*pers*: 0.85) and D306 for R685 (*pers*: 0.90) (see **Figure 3.C**). These three key anionic residues involved in the highest persistent arginine – salt-bridges (in _681_PRRAR_685_) was further envisaged to form a combined molecular entity (let’s call it the ‘anionic triad’) made up of discrete molecular components (anionic side chains). The anionic triad and the catalytic triad (D153, H194, S368) remained proximal and approachable (average centroid–to–centroid distance between side chain atoms: 13.16 Å, SD: 0.72) throughout the 300 ns trajectory (RR1_CoV-2_) with the display of correlated movements as if like an ordered pair (**Figure 4.A**). To analytically confirm the visually apparent correlated movement, the inter-triad angle (ε), defined as the planer internal angle subtended by the two position vectors originated from the Furin molecular centroid that connect the centroids of the two triads (**Figure 4.B**), was surveyed across the trajectory. The inter-triad angle was found to be strictly constrained (<ε>=45.1°, SD=4.7) having an apparently bi-modal distribution with two modes at ∼42.1° and ∼52.8°.

**Figure 3.**
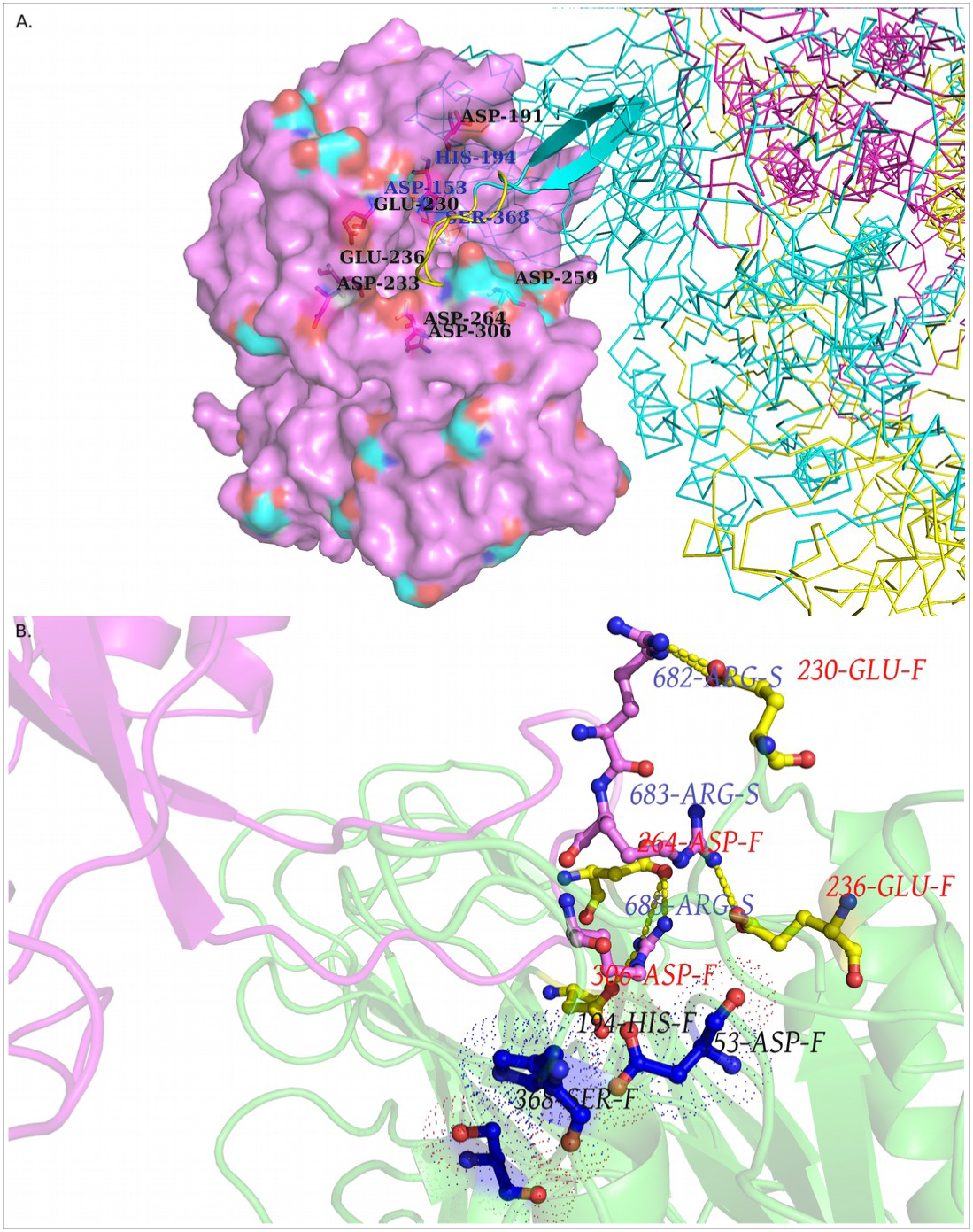
FLCS_Spike_ docked onto Furin and stabilized by Spike–Furin interfacial salt-bridges: as revealed from RR1_CoV2_. Panel **A** maps the docked FLCS_Spike_ (loop, highlighted in yellow, flanked at either end by beta-strands colored in cyan) at the Furin docking site in RR1_CoV2_ while the rest of the Spike that is visible in this closeup view is in ribbon (PyMol). A direct visual comparison can be made between Figure 2 portraying the proposed dockable surface groove (see corresponding main-text, section **3.4**) near the Furin catalytic triad and **Figure 3.A** showing the docked FLCS_Spike_ (as it occurred) at the very groove, surrounded by anionic residues amenable to form salt-bridges with the FLCS_Spike_ – arginines (R682, R683, R685). Panel **B** highlights the highest persistence Spike–Furin interfacial salt-bridges (thick yellow dashed lines) for the three arginines in the _681_PRRAR_685_ (FLCS_Spike_) along its 300 ns MD simulation trajectory (produced from RR1_CoV2_). Only a close up view of the interface is portrayed to highlight the key (highly persistent) salt-bridges (*pers*>0.85) sustaining the interface. Only the Furin – proximal part (FLCS_Spike_ with short flanking regions at either end) of the Spike (chain S) is shown in magenta cartoon (with R682, R683, R685 highlighted in ‘balls and sticks’) while the Furin chain (chain F) is drawn in green cartoon with its counter-ionic residues involved in the (aforementioned) key Spike–Furin interfacial salt-bridges highlighted in ‘balls and sticks’. Residues comprising the catalytic triad are presented in ‘balls and sticks’ colored in navy blue and highlighted by dot surface points. Figures were built in PyMol.

**Figure 4.**
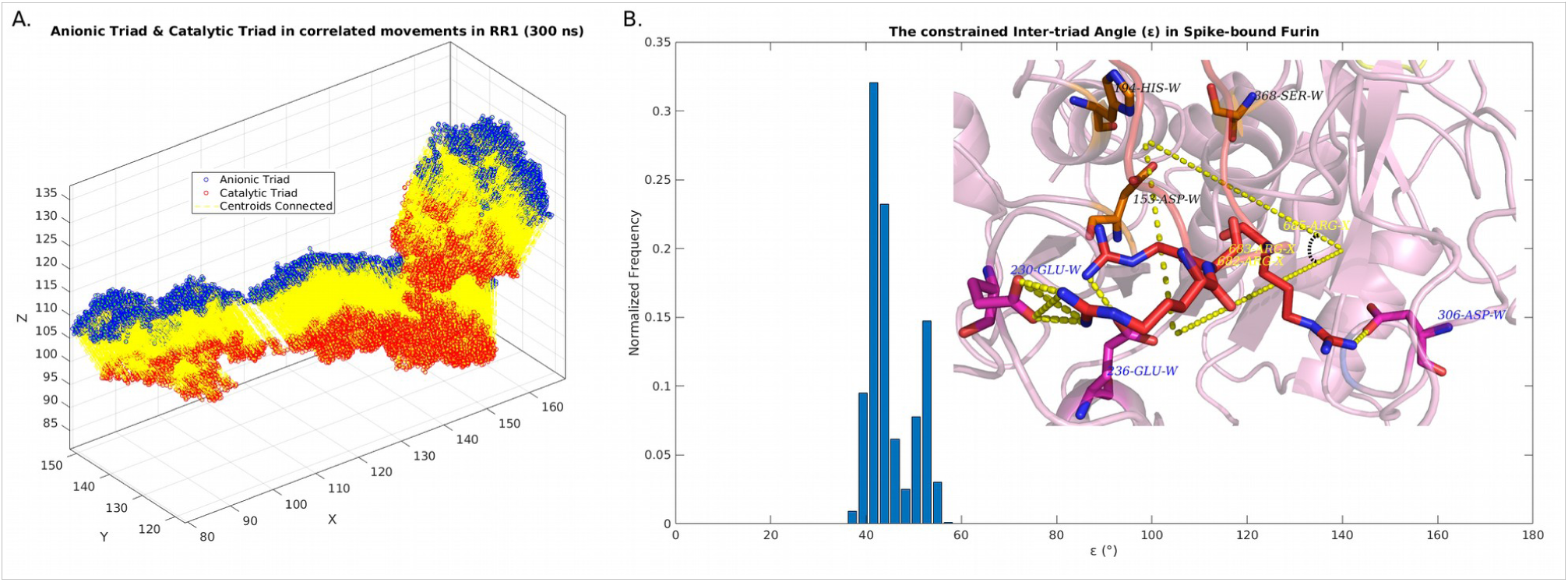
The anionic and the catalytic triad of the Spike – bound Furin in correlated movements throughout the 300 ns MD simulation trajectory (RR1).

Other interfacial salt-bridges barring those involving the three arginines in _681_PRRAR_685_ (FLCS_Spike_) was also surveyed in the same details. Among these, one salt-bridge, ‘E654_Spike_ ↔ R193_Furin_’, (see **Table 1**) was noticeable both in terms of its high persistence (*pers*: 0.65) and the opposite trend in the distribution of its charge centers (negative in Spike and positive in Furin, for a change) in contrast to the arginine – salt-bridges (in _681_PRRAR_685_). Other salt-bridges with brief/instantaneous occurrences (*pers<0.1*) could be considered ephemeral. The collective interplay of these fleeting salt-bridges triggers a ‘transient dynamics’ in disordered protein regions that is indispensable in retaining their flexibility [49] and is also pivotal towards imparting a critical behavior in associated disorder transitions among multiple self-similar fractal states [50]. The presence of transient salt-bridges (*pers*<0.1) in significant fractions (70.5% in the dynamic ensemble of RR1_CoV-2_, see **Table 1**) signals for relatively ordered metastable Furin-bound states of the FLCS_Spike_ which together retain enough flexibility (see **Video S1, Supplementary Materials**) to favor the Spike–S1/S2 proteolytic cleavage [33, 35].

#### 3.6.2. In ZR1_CoV-2_

As a cross-validation of the ‘salt-bridge hypothesis’, an independent guided docking was performed in Zdock using IRaPPA re-ranking (see section **2.3.1.2**, **Materials and Methods**) and the returned top ranked docked pose (ZR1_CoV-2_) was simulated in yet another independent MD simulation run for 100 ns (see section **2.4**, **Materials and Methods**). Following on, structural snapshots were sampled at 10 ps intervals (likewise to that of RR1_CoV-2_) leading to 10000 snapshots. Salt-bridges were then extracted from each snapshot from its interfacial contact map (see section **2.5.1**, **Materials and Methods**) and further sorted based on their dynamic persistence. A second sorted list, equivalent to that of RR1_CoV-2_ (**Table 1**), was procured for ZR1_CoV-2_ (**Table S5, Supplementary Materials**). Counter-ionic partners in most high persistence arginine – salt-bridges (in _681_PRRAR_685_) were preserved in both (dynamic) ensembles (RR1_CoV-2_, ZR1_CoV-2_) pairing either with the same (R682_Spike_ ↔ E230_Furin_, *pers*: 0.93, 0.99 respectively) or altered partners (R683_Spike_ ↔ E236_Furin_, *pers*: 0.85 in RR1_CoV-2_; R685_Spike_ ↔E236_Furin_, *pers*: 0.97 in ZR1_CoV-2_). Noticeably, 306-Asp (Furin) in RR1_CoV-2_ (R685_Spike_ ↔ D306_Furin_, *pers*: 0.90) was replaced by 259-Asp (Furin) in ZR1_CoV-2_ (R683_Spike_↔D259_Furin_, *pers*: 0.49). This suggests that there might be multiple plausible conformations (docked poses) mapping to unique (i.e., non-degenerate) yet befitting ionic bond network archetypes – all of which could enable the Spike–Furin binding. In fact to have such essential and nominal degrees of freedom generally in bio-molecular fitting (including self-fitting or folding) allows the system to breathe and is of no great surprise (at least) in protein science, often directed by satisfying optimized global physico-chemical constraints while retaining their structural degeneracy. The revelation of secondary and super-secondary structural motifs [113], packing motifs within native globular protein interiors [73], composite salt-bridge motifs within proteins and protein complexes [86] as well as the plausibility of the partly proven idea of alternative packing modes potentially leading to the same native protein fold (or a befitting hydrophobic core) [114, 115] – each and all are instances of the phenomena.

The Spike–Furin interfacial salt-bridges (in both RR1_CoV-2_ and ZR1_CoV-2_) generally varied in terms of their contact intensities (CI) (see section **2.5.2.2, Materials and Methods**) while the high persistence salt-bridges (formed near the Furin catalytic triad) were by and large, densely connected throughout their entire simulation runs (**Figure 5**), hitting appreciably high ACI values in most cases (**Table 1**, **Table S5**). Most high persistence salt-bridges also frequently retained maximally connected (CI=CI_max_=4) closed ionic bond (bipartite) motifs between bifurcated oppositely charged side chain groups in both subjects (**Figure 5, insets**).

**Figure 5.**
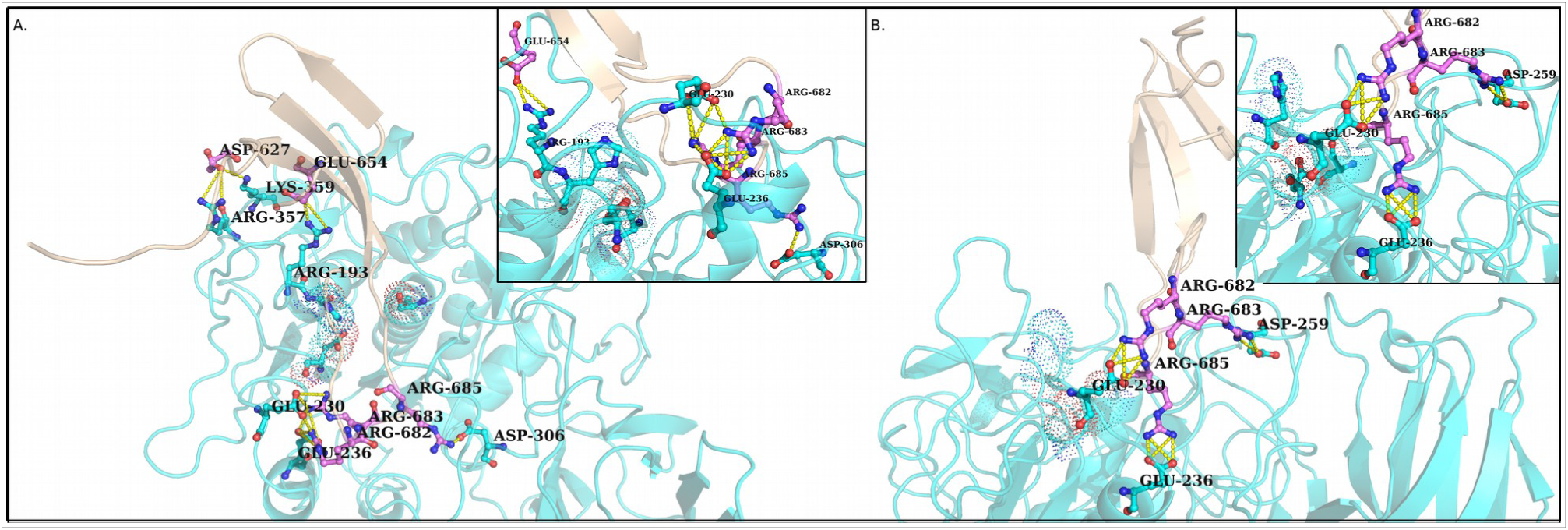
Densely connected composite ionic bond motifs in the persistent salt-bridges formed at the Spike–Furin interface. Panels A. and B. plots a representative snapshots of the interface (with a close-up view for the ionic bond motifs) randomly picked from the trajectories of RR1_CoV2_ and ZR1_CoV2_ respectively.

RR1_CoV-2_ and ZR1_CoV-2_ also had great resemblance in their frequency distribution profiles for the Spike–Furin interfacial salt-bridge persistence(s) (unweighted as well as weighted: see section **2.5.2.2, Materials and Methods**) [49] (**Figure S5, Supplementary Materials**). To quantify this resemblance, the entire theoretical range of persistence (*pers*) [0,1] was partitioned into 20 equally spaced bins, and for each ensemble, the normalized frequencies of salt-bridges falling within each persistence–bin (of bin-width: 0.05) were computed. A similar approach was adapted for weighted persistence (*wpers*) (see section **2.5.2.2, Materials and Methods)** which maps to a theoretical range of [0,4], equaling a bin-width of 0.2 for a 20-bin model. The Pearson Correlation between these obtained normalized frequencies from the two ensembles (RR1_CoV-2_, ZR1_CoV-2_) was found to be 0.97 for persistence (**Figure S5.A)** and 0.93 for weighted persistence (**Figure S5.B)** (*P-value<0.00001 for both which is significant at 99.9% level*). The same correlation (Pearson’s) was found to be 0.66 (*P-value: 0.001445, significant at 99.9% level*) between the frequency distribution profiles (RR1_CoV-2_, ZR1_CoV-2_) which are plotted for average contact intensities (ACI) of ionic bonds formed at the Spike–Furin interface (**Figure S5.A, inner set, Supplementary Materials**).

#### 3.6.3. In ZR1_CoV_, the baseline

As introduced in section **2.3.1.3** (**Materials and Methods**), ZR1_CoV_ (the representative Spike–Furin interaction in SARS-CoV, 2002/2003) served as the baseline for the Spike–Furin interaction in SARS-CoV-2. As has been already discussed (see section **2.2**, **Materials and Methods**), the FLCS_Spike_ patch in ZR1_CoV_ (_664_SLLRSTS_670_, originally missing in 7AKJ) is much shorter than its homologous missing patch in 6XR8 (_677_QTNS**PRRAR**SVA_689_) mapping to their corresponding degrees of disorder (higher in the later). The relative composition of the two patches and their pairwise alignments (**Figure S1, Supplementary Materials**) further support the observation that indeed a lesser degree of disorder is expected for the concerned patch in SARS-CoV than that in SARS-CoV-2. Most notably, the third arginine (R685) of _681_PRRAR_685_ in CoV-2 FLCS_Spike_ is evolutionarily conserved (as R667) also in CoV FLCS_Spike_ (see section **2.2**) (**Figure S1, Supplementary Materials**). A closer look into the pairwise alignments of the two sequences (**Figure S1, Supplementary Materials**) also reveals the strategic insertion of a dipeptide unit (_681_PR_682_) followed by two non-synonymous replacements (L665→R683, L666→A684) in the disordered activation loop (see **Introduction**) of the CoV-2 Spike (6XR8) with respect to its ancestral homologous Spike in CoV (7AKJ). With this background, when we had a good look at the MD simulation trajectories of ZR1_CoV_ we found something very insightful. The conserved arginine (R667) in 7AKJ was found to cover much space in ZR1_CoV_ throughout its entire dynamic trajectory (100 ns) together with an accompanying nearby lysine (K672), feeling up for the absence of the other arginines (in reference to _681_PRRAR_685_, CoV-2) leading to the formation of a homologous dynamically persistent network of interfacial salt-bridges in CoV. The conserved arginine, R667 alone seemed to engage as many as three counter-ionic Furin side chains (E230, E257, D258) forming two high persistent (*pers*: 0.62, 0.95) and one moderately persistent (*pers*: 0.28) salt-bridges (see **Table S6, Supplementary Materials**), while the neighboring lysine (K672) was found in pair with D258 (Furin) with a persistence of 0.58. Notably, D258 among the Furin anionic residues, shared persistent salt-bridges simultaneously with K672 and R667 (see **Table S6, Supplementary Materials**). In contrast to the _681_PRRAR_685_ (CoV-2 Spike), here, in context to _664_SLLRSTS_670_ (CoV Spike), the absence of the long and electrostatically repelling neighboring arginine side chains offers R667 the physical space to remain substantially flexible to be simultaneously involved in multiple high persistence salt-bridges. The formation of these interfacial salt-bridges are favored by the proximal looping of the flanking lysine (see **Figure S6, Supplementary Materials**). The two non-adjacent basic residues (R667, K672) together serves to sustain the homologous Spike–Furin interface in CoV, also, by the formation of several dynamically persistent ionic bonds. To that end, if the emergence of the ‘PRRAR’ motif (in CoV-2 Spike) is to be considered a solution that is optimized for the most efficient Spike–S1/S2 cleavage at the Spike–Furin interface, the interplay of R667 and K672 in context to the homologous FLCS_Spike_ in CoV appears to be analogous to the event of structural relaxation in mutant protein cores. In other words, the way R667 and K672 cover up the physical space in SARS-CoV to sustain the Furin and catalyze the Spike–S1/S2 cleavage appear to resemble with the collective conformational readjustments of neighboring residues, filling up for packing defects and/or cavities/holes introduced upon hydrophobic substitutions/truncation in native protein core(s) [116, 117]. Having said that, the total number of non-redundant Spike–Furin interfacial salt-bridges were found to be literally doubled by the incorporation of the additional arginines (P**RR**AR) in CoV-2 (17: RR1_CoV-2_, 20: ZR1_CoV-2_) compared to the baseline (9: ZR1_CoV_) in CoV. Also, the ionic bond networks seemed to be certainly more dense and intense in CoV-2 with a CCI (see section **2.5.2.2, Materials and Methods**) of 7.65, 10.16 in RR1_CoV-2_, ZR1_CoV-2_ compared to 5.48 in CoV (ZR1_CoV_). These are signatures of the proposed optimization characteristic of ‘gain of function’ mutational studies (see **Introduction**).

### 3.7. Enthalpy - entropy compensation involved in the Spike–Furin interaction

The ‘salt-bridge hypothesis’ (see section **3.5**) was proposed based on the intuition that the disordered high-entropy FLCS_Spike_ loop must get jammed into a restricted set of ‘allowed’ conformations into Furin that are favorable for the Spike–S1/S2 cleavage. Such a conformational selection should necessarily accompany electrostatic stabilization of the highly localized positive charge cloud on _681_PRRAR_685_ (FLCS_Spike_). The Spike–Furin interaction thus implicitly speaks in favor of a ‘*disorder-to-order transition*’ that needs an enthalpic source to compensate for the high entropic cost (loss) intrinsic to the supposed transition. One prime source for such enthalpic compensation are salt-bridges for they impart local rigidity in proteins by jamming conformations [86]. To that end, the ‘salt-bridge hypothesis’ was only found more plausible by the detection of a potentially dockable surface groove to fit in the FLCS_Spike_ near the Furin catalytic triad (see section **3.5**), surrounded by exposed anionic residues that seemingly have the potential to the form salt-bridges with the FLCS_Spike_ – arginines (in _681_PRRAR_685_). All structural dynamics analyses (see section **3.6**) unanimously and collectively reveal that the FLCS_Spike_ fits nicely and stably into the proposed dockable groove, stabilized by the formation and sustenance of dynamically interchangeable interfacial salt-bridges (validated in RR1_CoV-2_, and, cross-validated in ZR1_CoV-2_). Together these results speak in favor of an ‘enthalpy entropy compensation’ intrinsic to the transition from the disordered (free) to the relatively ordered (bound) state of the SARS-CoV-2 FLCS_Spike_. To further confirm this thermodynamic phenomenon, we computed actual structure – based all atom thermodynamic parameters by FoldX (see section **2.6**, **Materials and Methods**) for the respective MD simulation trajectories pertaining to RR1_CoV-2_, ZR1_CoV-2_ (300 ns, 100 ns) and compared the relevant transition enthalpic (ΔH_vdw_, ΔH_elec_) and entropic (ΔS_mc_, ΔS_sc_) terms associated with the Spike–Furin binding / complexation. Both molecules in their integral forms were considered for the FoldX energy calculations. To set up an appropriate baseline, ZR1_CoV_ was also included in the calculations and comparison. All relevant transition enthalpic and transition entropic terms individually as well as collectively had retained a counter trend (ΔH_vdw/elec_<0, ΔS_mc/sc_>0) throughout the entire trajectories of all subjects, RR1_CoV-2_, ZR1_CoV-2_, as well as the baseline, ZR1_CoV_ (**Table 2**, **Figure 6**). This is suggestive of enthalpy – entropy compensations accompanying both Spike–Furin binding events (in CoV-2, CoV). However, as is reflected from the relative magnitudes of the ΔH, ΔS terms (**Table 2, Figure 6**), the binding in CoV-2 (RR1_CoV-2_, ZR1_CoV-2_) is attributed with higher entropic costs of the event and therefore with a concomitant higher degree of enthalpic compensation than that in CoV (ZR1_CoV_).

**Figure 6.**
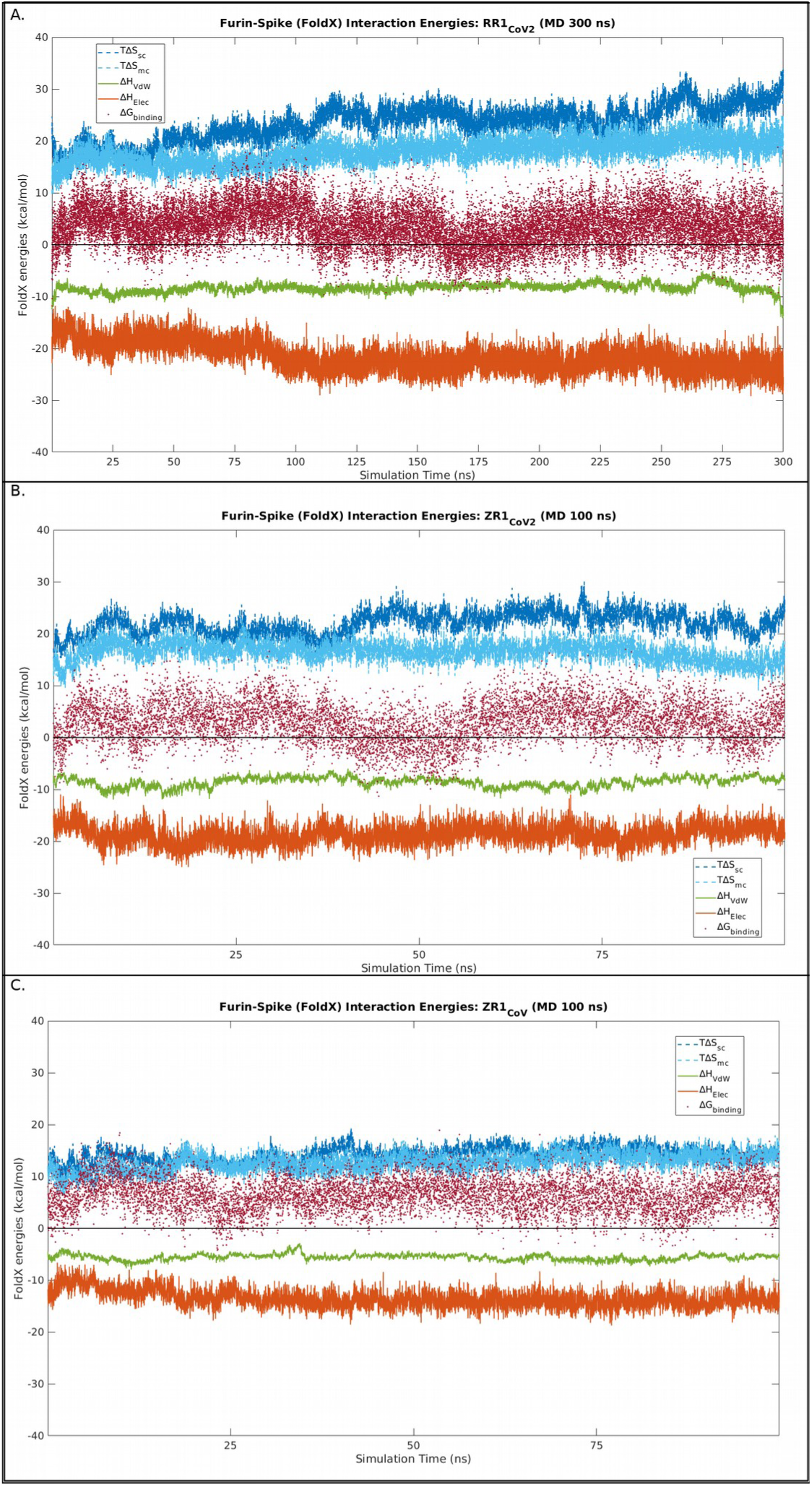
Interaction energy profiles (FoldX) for the top re-ranked Spike–Furin complexes (panl A. for RR1_CoV2_, panel B. for ZR1_CoV2_, panel C. for ZR1_CoV_) along their full MD simulation trajectories (300 ns, 100 ns, 300 ns). The different transition enthalpic (ΔH_vdw_, ΔH_elec_) and entropic (TΔS_mc_, TΔS_sc_) terms along with the net ΔG_binding_ are plotted in different colors as has been mentioned in the corresponding legend-boxes. All thermodynamic parameters are essentially energy terms and are plotted in the units of kcal mol^-1^.

**Table 2.** Structure-based equilibrium thermodynamic parameters (FoldX) obtained from the MD simulation trajectories of RR1_CoV2_, ZR1_CoV2_, ZR1_CoV_ representing the Spike–Furin binding in SARS-CoV2 (subject, cross-validation) and SARS-CoV (baseline) respectively. Time-series averages (in kcal mol^-1^) over the corresponding trajectories for each subject (RR1_CoV2_, ZR1_CoV2_, ZR1_CoV_) are given for each energy term (including enthalpic, entropic, free-energy terms) with their standard deviations given in parenthesis.

| | $\langle \Delta H_{\text{vdw}} \rangle$ | $\langle \Delta H_{\text{elec}} \rangle$ | $\langle \Delta S_{\text{mc}} \rangle$ | $\langle \Delta S_{\text{sc}} \rangle$ | $\langle \Delta G_{\text{binding}} \rangle$ |
| --- | --- | --- | --- | --- | --- |
| RR1<sub>CoV2</sub> | -21.722<br>(2.669) | -8.296<br>(0.871) | 17.890<br>(2.276) | 23.386<br>(3.432) | 3.687<br>(3.868) |
| ZR1<sub>CoV2</sub> | -18.748<br>(1.871) | -8.617<br>(0.956) | 16.498<br>(1.889) | 22.004<br>(2.243) | 3.043<br>(3.843) |
| ZR1<sub>CoV</sub> | -13.566<br>(1.695) | -5.556<br>(0.582) | 12.663<br>(1.746) | 14.218<br>(1.506) | 6.468<br>(3.135) |

As a second test, the individual main chain and side chain conformational entropies for the receptor (Spike) and the ligand (Furin) were also surveyed in RR1_CoV-2_, ZR1_CoV-2_, ZR1_CoV_ (throughout their respective trajectories) and compared between their unbound (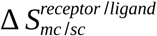) and bound (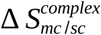) states. Entropic terms derived independently from both binding partners (Spike and Furin) in their unbound states (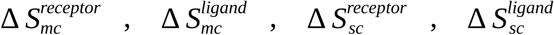; see section **2.6**, **Materials and Methods**) were found to be fairly stable over time (**Figure S7**, **Supplementary Materials**) – all of which get drastically reduced (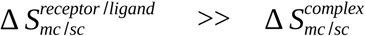 ; see **Table 3**) confirming the ‘entropy arrest’ (refer to section **2.6**, **Materials and Methods**) implicit to the Spike–Furin complexation in both systems (CoV-2, CoV). Though, the comparative transition entropy profiles and the time-averages were in the same range of values for both systems, literally all the surveyed terms had a rise of about 3-5% in terms of their average trends from the former to the later complex (**Table 3**). Perhaps with no great surprise, the most prominent rise (CoV→CoV-2) was found for the side chain conformational entropies (5.7%) of the receptor molecule ( 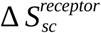, i.e., Spike) undergoing the transition, naturally for the sequence differences pertaining to the FLCS_Spike_ in both.

**Table 3.** Average entropic changes pertaining to the Spike–Furin interaction in RR1_CoV2_ and ZR1_CoV2_. The statistics is done over the entire 300 ns trajectory (RR1). Standard deviations given in parentheses. The subscripts in the ΔS terms refer to mc: main-chain, sc: side-chain.

|  | Receptor<br>(Spike) |  |  | Ligand<br>(Furin) |  |  | Complex<br>(Spike–Furin) |  |  |
| --- | --- | --- | --- | --- | --- | --- | --- | --- | --- |
|  | RR1<sub>CoV2</sub> | ZR1<sub>CoV2</sub> | ZR1<sub>CoV</sub> | RR1<sub>CoV2</sub> | ZR1<sub>CoV2</sub> | ZR1<sub>CoV</sub> | RR1<sub>CoV2</sub> | ZR1<sub>CoV2</sub> | ZR1<sub>CoV</sub> |
| $\langle \Delta S_{mc} \rangle$<br>(kcal<br>mol<sup>-1</sup>) | 6573.9<br>(21.8) | 6564.5<br>(23.8) | 6344.2<br>(23.5) | 2691.4<br>(16.1) | 2701.4<br>(14.8) | 2613.6<br>(14.3) | 17.9<br>(2.3) | 16.5<br>(1.9) | 12.7<br>(1.8) |
| $\langle \Delta S_{sc} \rangle$<br>(kcal<br>mol<sup>-1</sup>) | 2207.2<br>(12.9) | 2185.1<br>(14.7) | 2077.4<br>(21.4) | 860.8<br>(6.5) | 846.8<br>(7.4) | 821.9<br>(9.1) | 23.4<br>(3.4) | 22.0<br>(2.2) | 14.2<br>(1.5) |

The binding free energy overall was mildly disfavored (i.e., ΔG_binding_ mildly positive) in both Spike–Furin binding events (**Table 2**) suggesting perhaps to the characteristic formation of metastable and multi-stable interfaces throughout the coronavirus lineage. A strict negative ΔG_binding_ was obtained in 17.1%, 21.5% of the time-frames in the CoV-2 trajectories: RR1_CoV-2_, ZR1_CoV-2_, while the same fraction was found to be merely 1.8% in ZR1_CoV_, the baseline (in CoV). The metastabilities (suggesting an ‘on-and-off’ mode of binding) appear to be of no great surprise and perhaps anticipated given that the Spike–Furin binding works like a preface to the cleavage of a desired peptide bond that seems to be favored upon the transient formation of certain energetically favorable intermediate conformations in the bound FLCS_Spike_. In fact, given that such disordered activation loops (like that of FLCS_Spike_) are known to serve as key structural and kinetic determinants of protease substrates [40], it would be worth exploring (via future studies) across other families of proteases (see **Introduction**) harboring such cleavage loops, as to whether the metastabilities also holds true in them. Apart from the revealed characteristic metastabilities, the comparative free energy values for the Spike–Furin binding were roughly twice as much in magnitude in CoV (<ΔG_binding_>=6.429 kcal mol^-1^, SD=3.094 : ZR1_CoV_) compared to those in CoV-2 (<ΔG_binding_>= 3.687, 3.043 kcal mol^-1^, SD=3.868, 3.843 : RR1_CoV-2_, ZR1_CoV-2_). The corresponding ΔΔG_binding_ values for RR1_CoV-2_, ZR1_CoV-2_ were found to be -2.742, -2.561 kcal mol^-1^ (see section **2.6**, **Materials and Methods**) – which speak directly in favor of a much more facilitated transition in the evolutionarily later event, signaling for the intended optimization (irrespective of whether *natural* or not) to have indeed occurred in SARS-CoV-2.

### 3.8. Using the Ramachandran Plot to probe the ‘disorder-to-order transition’ of the SARS-CoV-2 FLCS_Spike_ loop upon Furin binding

Finally, the paper takes the opportunity to demonstrate a novel use of the legendary Ramachandran Plot (RP) [51] in probing the ‘*disorder-to-order transition*’ of the SARS-CoV-2 FLCS_Spike_ loop upon Furin binding. The unbound disordered state was taken to be the ensemble of 500 ‘all-atom’ SARS-CoV-2 Spike atomic models (see **section 3.2**) built with its experimentally missing patches modeled with uniquely varied loop-conformations (see section **2.2**, **Materials and Methods**). On the other hand, 500 time-frames sampled at equal temporal interval of 600 ps from the 300 ns MD simulation trajectory of the Spike–Furin complex (simulated from the rank-1 docked pose in the ClusPro blind-docking) were assembled to represent the bound (presumably ordered) state. The Ramachandran backbone torsion angles (*Φ, ψ*) were then computed for each atomic model under each ensemble (bound, unbound) for all Spike residues and those pertaining to the FLCS_Spike_ loop were extracted and plotted (overlaid) in the RP (**Figure 7**). The overlaid distributions clearly show more scatter for (*Φ, ψ*) points in the unbound (**Figure 7.A**) compared to bound (**Figure 7.B**) states. In addition, a re-view of the RPs were felt necessarily important with a sense of contiguity for the connected pentapeptide sequence motif (-_681_PRRAR_685_-) embedded in the FLCS_Spike_ loop. Such a sense of contiguity is also essential in terms of backbone tracing in protein crystallography [118] and depicting secondary structural elements [119]. To that end, a simple line-drawing of the successively connected residues belonging to the -P_681_-R_682_-R_683_-A_684_-R_685_- pentapeptide motif was performed (**Figure S8, Supplementary Materials**), over and above the standard ‘scattered points representation’ of the RP.

**Figure 7.**
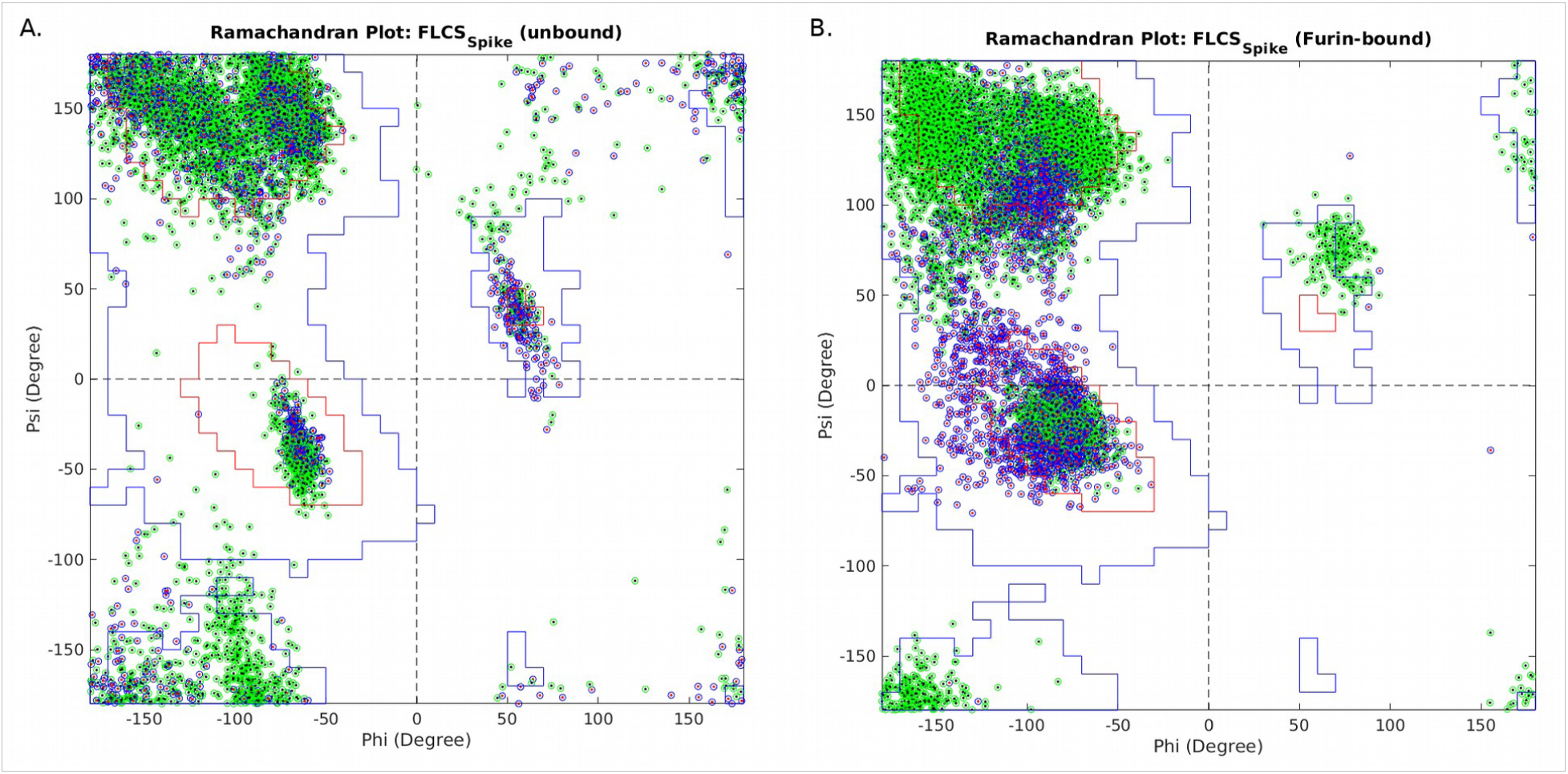
Overlaid Ramachrandran Plots for FLCS_Spike_ pertaining to (A) unbound and (B) Furin-bound Spike states. Each plot is overlaid with 100 atomic models belonging to the same state (unbound or bound). Within each individual plot (i.e., pertaining to each atomic model), the {Φ, Ψ}} points for residues in the FLCS_Spike_ loop is plotted as black dots encircled by green while those for residues belonging to the -P_681_-R_682_-R_683_-A_684_-R_685_- pentapeptide motif are highlighted by red dots encircled by blue.

**Figure 8.**
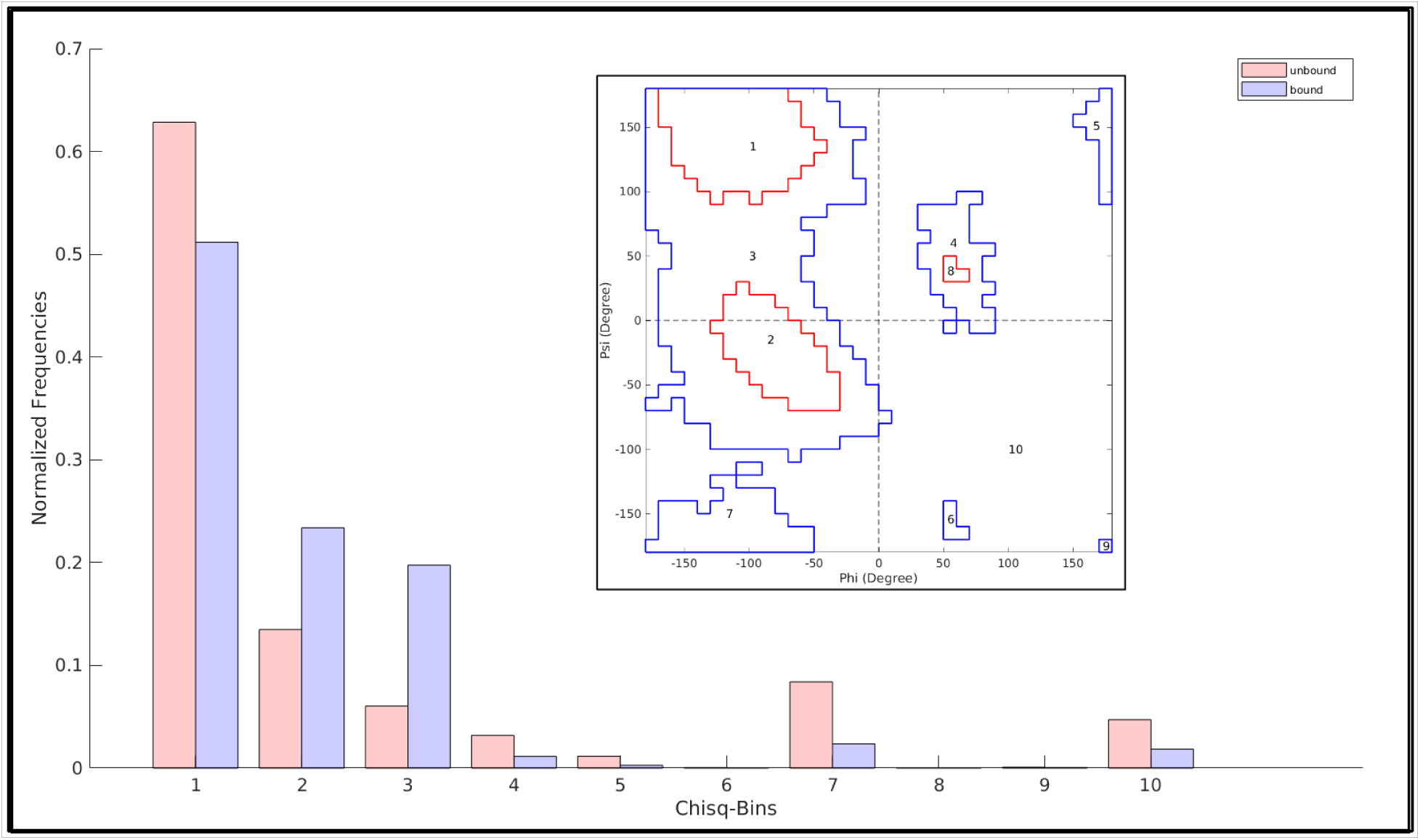
Distribution of differential counts obtained from binning of the RP for unbound and bound states of FLCS_Spike_. For binning details please refer to main-text. Bin-1, 2 & 8 represents Ramachandran allowed regions for β-sheets, Rα-helices, Lα-helices while 3-7, 9 maps to 6 disjoint partially allowed regions and bin-10 stands for the pulled-down disallowed region.

For the unbound state, the successive points clearly hovers around extreme ends of the RP resembling a highly multi modal distribution in terms of occupying different regions in RP indicating high structural conflicts or disorder. In comparison, the same successive points clearly gets shrunk into a constrained distribution for the Furin-bound state, directed to an extended ‘generously allowed’ region [120] of the RP. More interestingly, this extended region almost perfectly maps to the extended bridged territory of the originally proposed allowed regions [51] for beta-sheets and right handed alpha- (as well as 3_10_-) helices upon the relaxation of the bond-angle, tau (τ: N-Cα-C) [121]. Motivated by these very interesting observations, we further computed the τ – angle for the FLCS_Spike_ in both the ensembles (unbound and bound). The τ – angle in the Furin-bound state (time-averaged) was indeed found to be more relaxed (<τ_Furin-bound_>: 109.6°; SD^9^: 4.0°) and trending to its ideal value of 109.5° for a tetrahedral sp^3^ carbon [122], compared to its unbound state where the average value (<τ_unbound_>: 107.4°; SD: 2.3°) was somewhat left-shifted from its tetrahedral ideal value. When surveyed for the -_681_PRRAR_685_- pentapeptide motif, the two <τ> values were even more separated with similar ratio of their standard deviations (<τ_unbound_>: 107.9°; SD: 2.4°; <τ_Furin-bound_>: 110.8°; SD: 4.2°). In both cases (FLCS_Spike_, -_681_PRRAR_685_-), the standard deviations were 1.7 times more in the bound state (i.e., more relaxed τ – angles) than in the unbound state. Moreover, from the overlaid RPs plotted for the Furin-bound state, it appears highly likely that the flexible FLCS_Spike_ loop or at least a good part of the loop is in dynamic equilibrium with multiple short transient secondary structural elements (e.g., short helical turns, beta-strands etc.) in its bound state. Given the trends in {Φ, ψ}, rationalized by the comparatively relaxed τ – angle in the bound state, t}, rationalized by the comparatively relaxed τ – angle in the bound state, this appears especially relevant to the pentapeptide (-_681_PRRAR_685_-) motif (**Figure 7.B**).

Subsequent to the visual inspection and comparison, the RP derived parameters (see section **2.7**, **Materials and Methods**) were computed for the FLCS_Spike_ patch independently for each state (unbound and Furin-bound) to quantify the distribution of (*Φ, ψ*) points spanning across the two plots (**Figure 7.A & 7.B**) and to assess whether this difference is of any significance. From definition (see section **2.7**, **Materials and Methods**), smaller values of *|δδ_c_|δ* (say, <30°) statistically indicates that the consecutive residues are conformationally alike or close which effectively leads to a local structural coherence and relative structural order for the FLCS_Spike_, while a larger value suggest regular structural conflicts and consequently structural disorder. Thus, from definition, *|δδ_c_|δ* also gives an estimate of how much the FLCS_Spike_ is conformationally varied (or, in other words, distributed among varying structural conformations) on an average. Lesser values of *|δδ_c_|δ* indicate greater tendencies (on an average) of the FLCS_Spike_ to attain closely related structural conformations, while as the value increases, structural degeneracy [50] is manifested within the FLCS_Spike_.

All the RP-derived parameters (see section **2.7**, **Materials and Methods**) are spread descriptors of some sort and all of them unequivocally drop (i.e., shrink) in the bound state (**Table 4**) compared to the unbound state. The relative decrease from state-1 (Spike, unbound) to state-2 (Spike, Furin-bound) in this two-state transition is 32.5% in <δ>_3q_ and 55.7% in <δ>_9d_. This indicates that the distance (δ, defined in the *Φ-ψ* space) by which 90% of points are seperated in the two RPs (states) is increased more than 1.5 times in the unbound state, compared to the bound state. Together, the visual and the quantitative analyses clearly and directly portray (from actual structural dynamics data) the transition of the unbound disordered FLCS_Spike_ to a relatively ordered Furin-bound state in SARS-CoV-2 Spike.

**Table 4.** The RP-derived parameters for the FLCS_Spike_ patches pertaining to unbound and bound Spike states in SARS-CoV-2. The parameters in the header-cells (|δ_c_|, σ(δ_c_), <δ>_median_, <δ>_3q_, <δ>_9d_) are as defined in section 2.7, **Materials and Methods**.

| Spike States | $\vert \delta_c\vert $<br>($\sigma(\delta_c)$) | $\langle\delta\rangle_{\text{median}}$ | $\langle\delta\rangle_{3q}$ | $\langle\delta\rangle_{9d}$ |
| --- | --- | --- | --- | --- |
| State-1<br>(Unbound) | 138.5<br>(30.6) | 101.0 | 212.0 | 305.2 |
| State-2<br>(Bound) | 118.7<br>(26.2) | 94.7 | 160.5 | 196.0 |
| Relative increase<br>from state-1<br>to state-2<br>(%) | -16.7 | -6.7 | -32.1 | -55.7 |

Lastly, to render a statistical significance to the change in the obtained distributions of (*Φ, ψ*) points in the RP associated with the two state transition (unbound → bound) of the FLCS_Spike_ a χ^2^ test was performed. Ten distinct bins corresponding to disjoint regions in the RP was considered. We adapted the Procheck [123] version of the RP to reproduce the Ramachandran (*Φ, ψ*) contours (**Figure 7**) and used the MATLAB inbuilt function ‘*inpolygon*’ for the frequency distribution of (*Φ, ψ*) points into these bins. For reasons of simplicity, the later-extended generously allowed regions [120, 123] of the RP were avoided. The 10 bins thus represented 3 allowed regions for regular secondary structural elements (β-sheets, Rα-helices, Lα-helics)^10^, 6 partially allowed regions (of largely varying areas) across the plot, and, the entire left-over disallowed region, pulled into the 10^th^ bin. The null hypothesis was taken to be ‘*no or little (i.e., insignificant) changes caused in the unbound (Φ, ψ) points (FLCS_Spike_) upon binding to Furin (i.e., Expected: unbound; Observed: bound)*’. The *χ^2^* (see section **2.8**, **Materials and Methods**) value obtained from the differential counts (**Figure 7**) of points (unbound → bound) in this 10- bin distribution (df^11^=9) was found to be 3650.32 which is ∼131 times higher than that of the upper-tail critical *χ^2^* value for df=9 at 99.9% level of significance (*χ^2^* = 27.88). Based on these numbers, the null hypothesis was rejected which should mean that the ‘unbound → bound’ change was indeed significant in the *FLCS_Spike_* in terms of their relative RP distributions even at the 99.9% level. In other words, the ‘*disorder→order transition*’ of the *FLCS_Spike_* upon binding to Furin was evident and unmistakable.

The RP has previously been used to probe transitions among α-helix, Π-helix and turns in context to the Phosphorylation of Smooth Muscle Myosin [124]. Having said that, the visual impact of simple line-drawing (to portray sequence contiguity) as well as the collective use of RP derived metrices, to the best of our knowledge and belief, together presents yet another novel use of the evergreen and multifaceted Ramachandran Plot.

## 4. Conclusion and Perspective

In parallel to the ongoing efforts to find a sustainable therapeutic solution to curb the coronavirus pandemic, debates are also ongoing regarding the origin of SARS-CoV-2. The current paradigm is that of an accidental ‘lab escape’ of SARS-CoV-2 giving rise to COVID-19 (from an ongoing ‘gain of function’ mutational studies) which has found great support from Genome comparison studies of late (see **Introduction**) revealing the sudden emergence of the _681_PRRAR_685_ motif in the SARS-CoV-2 Spike, absent in other related respiratory viruses. The strategic presence of such polybasic motifs in FLCS_Spike_ like flexible loops in coronavirus and other related respiratory viral lineages leads to local protein disorder [21], intrinsic to these activation loops and the one in SARS-CoV-2 is believed to play a key role in the drastic increase in viral host cell entry and transmissibility. To the very best of our knowledge, the current study is the first of its kind that entraps a ’*disorder-to-order* transition’ in the SARS-CoV-2 FLCS_Spike_ while it undergoes host Furin binding that is optimized for a more efficient proteolytic cleavage of its S1/S2 junction than that in SARS-CoV. The optimization and the consequent increase in proteolytic cleavage efficiency is unambiguous from all analyses performed (sections **3.5**-**3.8**) but is perhaps the most clear and direct from the fairly negative ΔΔG_binding_ values returned from the two events (section **3.7**). The study further reveals the key role of dynamically interchangeable, persistent salt-bridges at the Spike–Furin interface – which seem to be an evolutionarily conserved feature of the coronavirus lineage and is substantially enhanced in the case of SARS-CoV-2 due to the presence of the three arginines (R682, R683, R685) in the _681_PRRAR_685_ motif amid its FLCS_Spike_. The host Furin, orchestrated with a preponderance of exposed amenable anionic residues (E230, E236, D259, D264, D306) strategically positioned around its catalytic triad overwhelmingly favors polybasic disordered substrates like that of the _681_PRRAR_685_ motif (SARS-CoV-2) for binding, cleavage and consequent host cell entry of the virus (sections **3.5**-**3.6**). The resultant Spike–Furin interfacial salt-bridges not only serves as a prominent enthalpy source for the process (compensating for the entropic loss of the FLCS_Spike_ undergoing ‘disorder-to-order transition’) but also favors the system to retain its characteristic metastabilities favorable for proteolytic cleavages targeted at flexible protein loops (section **3.7**). The current study also helps to open up new research avenues across other related protease families harboring such cleavage loops, as to whether this revealed metastabilities also holds true in them. The findings are perfectly consistent with the established theories of salt-bridge dynamics in context to IDPs serving to retain their characteristic structural plasticity by the continuous triggering of phase transitions among their self-similar disordered states [49, 50]. Further, from the combined results of salt-bridge and thermodynamic analyses (see section **3.6, 3.7**) it strongly appears that the Furin cleavage seeks opportunities for transient formation of favorable intermediate conformations in the bound FLCS_Spike_ to make the final unfailing strike on the desired peptide bond. The probabilities to have this successful strike is naturally far greater in SARS-CoV-2 (than in SARS-CoV) since it has the more intense salt-bridge networks formed and sustained in its Spike–Furin interface (for the presence of the _681_PRRAR_685_ – arginines). These findings further rationalize the substantially greater extent of cleavage (59.6%) of the SARS-CoV-2 Spike (into its S1/S2 products) in the wild-type virion than in its ΔPRRA mutant (14.5%) [33]. In conclusion, over and above offering a novel perspective into the coronavirus molecular evolution, the study also makes the SARS-CoV-2 Spike–Furin interaction mechanistically insightful adding new dimensions to the existing theories of proteolytic cleavages *per se*.

## Supporting information

The Supplementary Materials File (PDF) contains 8 Supplementary Figures, 6 Supplementary Tables & one Supplementary Video

## Acknowledgment and Funding

We convey our sincerest gratitude to Prof. Raghavan Varadarajan, Molecular Biophysics Unit, IISc, Bangalore, India and Prof. Dhananjay Bhattacharyya, Computational Science Division, Saha Institute of Nuclear Physics, Kolkata, India (retired) for their constant motivations and helpful discussions during the work. The project was self-funded.

## Author’s Contributions

Conceptualization: SB; Design: SB, AB; Setting up and performing the MD simulations: SR; Formal analysis and investigation: SB, AB, SR, PG; Methodology and validation: SB, AB, SR; Literature review: PG, SR, SB; Writing—original draft preparation: SB, AB; writing—review and editing SB, AB, PG, SR. All authors have read and approved the final manuscript.

## Supplementary Information

Enclosed, please find the Supplementary Materials File (in PDF format) containing eight Supplementary Figures, six Supplementary Tables and one Supplementary Video.

## Footnotes

1 ORF: Open Reading Frame

2 PDB: Protein Data Bank

3 RMSD: Root mean square deviations

4 df: degree of freedom

5 QM/MM: Quantum Mechanics / Molecular Mechanics

6 AI: Artificial Intelligence

7 FPR: False Positive Rate

8 SD: Standard Deviation

9 SD: Standard Deviation

10 R: Right handed; L: Left handed

11 df: Degree of Freedom

