## Supplementary material for "Capturing a crucial ‘disorder-to-order transition’ at the heart of the coronavirus molecular pathology – triggered by highly persistent, interchangeable salt-bridges": The Supplementary Materials File (PDF) contains 8 Supplementary Figures, 6 Supplementary Tables & one Supplementary Video

by

*Sourav Roy, Prithwi Ghosh, Abhirup Bandyopadhyay, Sankar Basu[*]*

**Table S1. Dataset of representative Coronavirus (CoV/CoV-2) Spike experimental (cryo-EM) structures with a resolution not worse than 3 Å, extracted from the PDB** (dated. 31st December, 2021).

| PDB ID | Resolution (Å) | Description (TITLE) | CoV/CoV-2 |
|---|---|---|---|
| 6m15 | 2.38 | Cryo-Em Structures Of Hku2 Spike Glycoproteins | CoV |
| 6m16 | 2.83 | Cryo-Em Structures Of Sads-Cov Spike Glycoproteins | CoV |
| 6nzk | 2.80 | Structural Basis For Human Corona Virus Attachment To Sialic Acid Receptors | CoV |
| 6ohw | 2.90 | Structural Basis For Human Corona Virus Attachment To Sialic Acid Receptors. Apo-Hcov-Oc43 S | CoV |
| 6q04 | 2.50 | Mers-Cov S Structure In Complex With 5-N-Acetyl Neuraminic Acid | CoV |
| 6q05 | 2.80 | Mers-Cov S Structure In Complex With Sialyl-Lewisx | CoV |
| 6q06 | 2.70 | Mers-Cov S Structure In Complex With 2,3-Rialyl-N-Acetyl-Lactosamine | CoV |
| 6q07 | 2.90 | Mers-Cov S Structure In Complex With 2,6-Rialyl-N-Acetyl-Lactosamine | CoV |
| 7bbh | 2.90 | Structure Of Coronavirus Spike From Smuggled Guangdong Pangolin | CoV |
| 7cn4 | 2.93 | Cryo-Em Structure Of Bat Ratg13 Spike Glycoprotein | CoV |
| 7cn8 | 2.50 | Cryo-Em Structure Of Pcov_Gx Spike Glycoprotein | CoV |
| 7m5e | 2.50 | Mers-Cov S Bound To The Broadly Neutralizing B6 Fab Fragment (C3 Refinement) | CoV |
| 6vxx | 2.80 | Structure Of The Sars-Cov-2 Spike Glycoprotein (CLOSED State) | CoV-2 |
| 6x29 | 2.70 | Sars-Cov-2 Rs2d Down State Spike Protein Trimer | CoV-2 |
| 6x79 | 2.90 | Prefusion Sars-Cov-2 S Ectodomain Trimer Covalently Stabilized In The<br>Closed Conformation | CoV-2 |
| 6xlu | 2.40 | Structure Of Sars-Cov-2 Spike At Ph 4.0 | CoV-2 |
| 6xm0 | 2.70 | Consensus Structure Of Sars-Cov-2 Spike At Ph 5.5 | CoV-2 |
| 6xm3 | 2.90 | Structure Of Sars-Cov-2 Spike At Ph 5.5, Single Rbd Up, Conformation 1 | CoV-2 |
| 6xm4 | 2.90 | Structure Of Sars-Cov-2 Spike At Ph 5.5, Single Rbd Up, Conformation 2 | CoV-2 |

| | | | |
|---|---|---|---|
| 6xr8 | 2.90 | Distinct Conformational States Of Sars-Cov-2 Spike Protein | CoV-2 |
| 6xra | 3.00 | Distinct Conformational States Of Sars-Cov-2 Spike Protein | CoV-2 |
| 6zb5 | 2.85 | Sars Cov-2 Spike Protein, Closed Conformation, C3 Symmetry | CoV-2 |
| 6zge | 2.60 | Uncleavable Spike Protein Of Sars-Cov-2 In Closed Conformation | CoV-2 |
| 6zgi | 2.90 | Furin Cleaved Spike Protein Of Sars-Cov-2 In Closed Conformation | CoV-2 |
| 6zow | 3.00 | Sars-Cov-2 Spike In Prefusion State | CoV-2 |
| 6zox | 3.00 | Structure Of Disulphide-Rtabilized Sars-Cov-2 Spike Protein Trimer (X2 Disulphide-Bond Mutant, G413c, V987c, Single Arg S1/S2 Cleavage Site) | CoV-2 |
| 6zp0 | 3.00 | Structure Of Sars-Cov-2 Spike Protein Trimer (SINGLE Arg S1/S2 Cleavage Site) In Closed State | CoV-2 |
| 7a4n | 2.75 | Cryo-Em Structure Of A Prefusion Stabilized Sars-Cov-2 Spike (D614N, R682s, R685g, A892p, A942p And V987p)(S-Closed Trimer) | CoV-2 |
| 7ad1 | 2.92 | Cryo-Em Structure Of A Prefusion Stabilized Sars-Cov-2 Spike (D614N, R682s, R685g, A892p, A942p And V987p)(One Up Trimer) | CoV-2 |
| 7ddd | 3.00 | Sars-CoV-2 S Protein At Close State | CoV-2 |
| 7df3 | 2.70 | Sars-Cov-2 S Trimer, S-Closed | CoV-2 |
| 7dwy | 2.70 | S Protein Of Sars-Cov-2 In The Locked Conformation | CoV-2 |
| 7e7b | 2.60 | Cryo-Em Structure Of The Sars-Cov-2 Furin Site Mutant S-Trimer From A Subunit Vaccine Candidate | CoV-2 |
| 7jwy | 2.50 | Structure Of Sars-Cov-2 Spike At Ph 4.5 | CoV-2 |
| 7kdk | 2.80 | Sars-Cov-2 D614g 3 Rbd Down Spike Protein Trimer Without The P986-P987 Stabilizing Mutations (S-GSAS-D614G) | CoV-2 |
| 7kdl | 2.96 | Sars-Cov-2 D614g 1-Rbd Up Spike Protein Trimer Without The P986-P987 Stabilizing Mutations (S-GSAS-D614G) | CoV-2 |
| 7lwk | 2.92 | Mink Cluster 5-Associated Sars-Cov-2 Spike Protein (S-GSAS-D614G-DELFV) In The 3-Rbd Down Conformation | CoV-2 |
| 7lwl | 2.84 | Mink Cluster 5-Associated Sars-Cov-2 Spike Protein (S-GSAS-D614G-DELFV) In The 3-Rbd Down Conformation | CoV-2 |
| 7lwm | 2.83 | Mink Cluster 5-Associated Sars-Cov-2 Spike Protein (S-GSAS-D614G-DELFV) In The 1-Rbd Up Conformation | CoV-2 |
| 7lwn | 2.94 | Mink Cluster 5-Associated Sars-Cov-2 Spike Protein | CoV-2 |

| | | | |
|---|---|---|---|
| | | (S-GSAS-D614G-DELFV) In The 1-Rbd Up Conformation | |
| 7lwo | 2.85 | Mink Cluster 5-Associated Sars-Cov-2 Spike Protein (S-GSAS-D614G-DELFV) In The 1-Rbd Up Conformation | CoV-2 |
| 7lww | 3.00 | Triple Mutant (K417N-E484K-N501Y) Sars-Cov-2 Spike Protein In The 1-Rbd-Up Conformation (S-GSAS-D614G-K417N-E484K-N501Y) | CoV-2 |
| 7mjg | 2.81 | Cryo-Em Structure Of The Sars-Cov-2 N501y Mutant Spike Protein Ectodomain | CoV-2 |
| 7mtc | 2.60 | Structure Of Freshly Purified Sars-Cov-2 S2p Spike At Ph 7.4 | CoV-2 |

**Figures S1. Pairwise alignments of the FLCS$_{Spike}$ sequences extracted from 7AKJ (SARS-CoV Spike) and 6XR8 (SARS-CoV-2 Spike).**

```
>7akj_EM   ASYHTV-------QKSIVAYTMSLGADSSIAYSNNTIAIPTNFSISITTEVMPVSM       720
>7akj_PM   ASYHTVSLLRSTSQKSIVAYTMSLGADSSIAYSNNTIAIPTNFSISITTEVMPVSM       720

>6xr8_EM   ASYQT------------SQSIIAYTMSLGAENSVAYSNNSIAIPTNFTISVTTEILPVSM   720
>6xr8_PM   ASYQTQTNSPRRARSVASQSIIAYTMSLGAENSVAYSNNSIAIPTNFTISVTTEILPVSM   720

>7akj_PM   ASYHTVS----LLRSTSQKSIVAYTMSLGADSSIAYSNNTIAIPTNFSISITTEVMPVSM   720
>6xr8_PM   ASYQTQTNSPRRARSVASQSIIAYTMSLGAENSVAYSNNSIAIPTNFTISVTTEILPVSM   720
```

```
PM: Proteomic (Sequence) Data
EM: Electron Microscopic (Structural) Data

7akj: SARS-CoV  (2003) Spike
6xr8: SARS-CoV2 (2019) Spike
```

**Figures S2. Multiple Sequence alignment (MSA) of the coronavirus FLCS$_{Spike}$ sequences with the unique pentapeptide activation loops highlighted (colored differentially).** MSAs were performed by MUSCLE [1] and cross-validated by CLUSTAL OMEGA [2].

```
6m15    NPLGDGFCADLLSNVVV---------RRMTFEKHDTTY-------VAPVTNERFTELPLD   569
6m16    NPLGDGFCADLLGNVAV---------RRMTFEKHDTTY-------VAPVTNERYTEMPLD   571
6nzk    LTVGSGYCVDYSKNGGSGGA----ITTGYRFTNFEPFTVNSVNDSLEPVGGLYEIQIPSE   805
6ohw    LTVGSGYCVDYSKNGGSGGA----ITTGYRFTNFEPFTVNSVNDSLEPVGGLYEIQIPSE   805
7bbh    IPIGAGICASYQTNTNS-----RSVSSQ---AII-AYTM-SLGAENSVAYANNSIAIPTN   709
7cn8    IPVGAGICASYHSMS-----SLRSVNQR---SII-AYTM-SLGAENSVAYSNNSIAIPTN   711
6zox    IPIGAGICASYQTQTNS-R----SVASQ---SII-AYTM-SLGAENSVAYSNNSIAIPTN   704
6zp0    IPIGAGICASYQTQTNS-R----SVASQ---SII-AYTM-SLGAENSVAYSNNSIAIPTN   704
6zge    IPIGAGICASYQTQTNS-PSRASSVASQ---SII-AYTM-SLGAENSVAYSNNSIAIPTN   748
6q04    LPLGQSLCALPDTPSTLTPASVGSVPGEMRLASI-AFNH-PIQV-DQLNSSYFKLSIPTN   799
6q05    LPLGQSLCALPDTPSTLTPASVGSVPGEMRLASI-AFNH-PIQV-DQLNSSYFKLSIPTN   799
6q06    LPLGQSLCALPDTPSTLTPASVGSVPGEMRLASI-AFNH-PIQV-DQLNSSYFKLSIPTN   799
6q07    LPLGQSLCALPDTPSTLTPASVGSVPGEMRLASI-AFNH-PIQV-DQLNSSYFKLSIPTN   799
7m5e    LPLGQSLCALPDTPSTLTPASVGSVPGEMRLASI-AFNH-PIQV-DQLNSSYFKLSIPTN   799
6vxx    IPIGAGICASYQTQTNS-PSGAGSVASQ---SII-AYTM-SLGAENSVAYSNNSIAIPTN   736
7mtc    IPIGAGICASYQTQTNS-PSGAGSVASQ---SII-AYTM-SLGAENSVAYSNNSIAIPTN   736
6x79    IPIGAGICASYQTQTNS-PSGAGSVASQ---SII-AYTM-SLGAENSVAYSNNSIAIPTN   736
7a4n    IPIGAGICASYQTQTNS-PSRAGSVASQ---SII-AYTM-SLGAENSVAYSNNSIAIPTN   717
7ad1    IPIGAGICASYQTQTNS-PSRAGSVASQ---SII-AYTM-SLGAENSVAYSNNSIAIPTN   717
7lwk    IPIGAGICASYQTQTNS-PGSASSVASQ---SVI-AYTM-SLGAENSVAYSNNSIAIPTN   715
7lwl    IPIGAGICASYQTQTNS-PGSASSVASQ---SVI-AYTM-SLGAENSVAYSNNSIAIPTN   715
7lwm    IPIGAGICASYQTQTNS-PGSASSVASQ---SVI-AYTM-SLGAENSVAYSNNSIAIPTN   715
7lwn    IPIGAGICASYQTQTNS-PGSASSVASQ---SVI-AYTM-SLGAENSVAYSNNSIAIPTN   715
7lwo    IPIGAGICASYQTQTNS-PGSASSVASQ---SVI-AYTM-SLGAENSVAYSNNSIAIPTN   715
7kdk    IPIGAGICASYQTQTNS-PGSASSVASQ---SII-AYTM-SLGAENSVAYSNNSIAIPTN   717
7kdl    IPIGAGICASYQTQTNS-PGSASSVASQ---SII-AYTM-SLGAENSVAYSNNSIAIPTN   717
7lww    IPIGAGICASYQTQTNS-PGSASSVASQ---SII-AYTM-SLGAENSVAYSNNSIAIPTN   717
7mjg    IPIGAGICASYQTQTNS-PGSASSVASQ---SII-AYTM-SLGAENSVAYSNNSIAIPTN   717
6x29    IPIGAGICASYQTQTNS-PGSASSVASQ---SII-AYTM-SLGAENSVAYSNNSIAIPTN   702
7df3    IPIGAGICASYQTQTNS-PGSASSVASQ---SII-AYTM-SLGAENSVAYSNNSIAIPTN   717
7ddd    IPIGAGICASYQTQTNS-PGSASSVASQ---SII-AYTM-SLGAENSVAYSNNSIAIPTN   717
6xlu    IPIGAGICASYQTQTNS-PGSASSVASQ---SII-AYTM-SLGAENSVAYSNNSIAIPTN   704
6xm0    IPIGAGICASYQTQTNS-PGSASSVASQ---SII-AYTM-SLGAENSVAYSNNSIAIPTN   704
6xm3    IPIGAGICASYQTQTNS-PGSASSVASQ---SII-AYTM-SLGAENSVAYSNNSIAIPTN   704
6xm4    IPIGAGICASYQTQTNS-PGSASSVASQ---SII-AYTM-SLGAENSVAYSNNSIAIPTN   704
6zow    IPIGAGICASYQTQTNS-PGSASSVASQ---SII-AYTM-SLGAENSVAYSNNSIAIPTN   717
7jwy    IPIGAGICASYQTQTNS-PGSASSVASQ---SII-AYTM-SLGAENSVAYSNNSIAIPTN   704
6zb5    IPIGAGICASYQTQTNS-PRRAASVASQ---SII-AYTM-SLGAENSVAYSNNSIAIPTN   717
6xr8    IPIGAGICASYQTQTNS-PRRARSVASQ---SII-AYTM-SLGAENSVAYSNNSIAIPTN   717
6xra    IPIGAGICASYQTQTNS-PRRARSVASQ---SII-AYTM-SLGAENSVAYSNNSIAIPTN   717
6zgi    IPIGAGICASYQTQTNS-PRRARSVASQ---SII-AYTM-SLGAENSVAYSNNSIAIPTN   748
7dwy    IPIGAGICASYQTQTNS-PRRARSVASQ---SII-AYTM-SLGAENSVAYSNNSIAIPTN   717
          :* *.                                               :* :
```

**Figure S3. Cluspro 2.0 – returned docked poses for a single run of 'ensemble (blind) docking' of Furin (ligand) and one of the SARS-CoV-2 Spike models (receptor) sampled from the CoV-2 $FLCS_{Spike}$ disordered ensemble.**

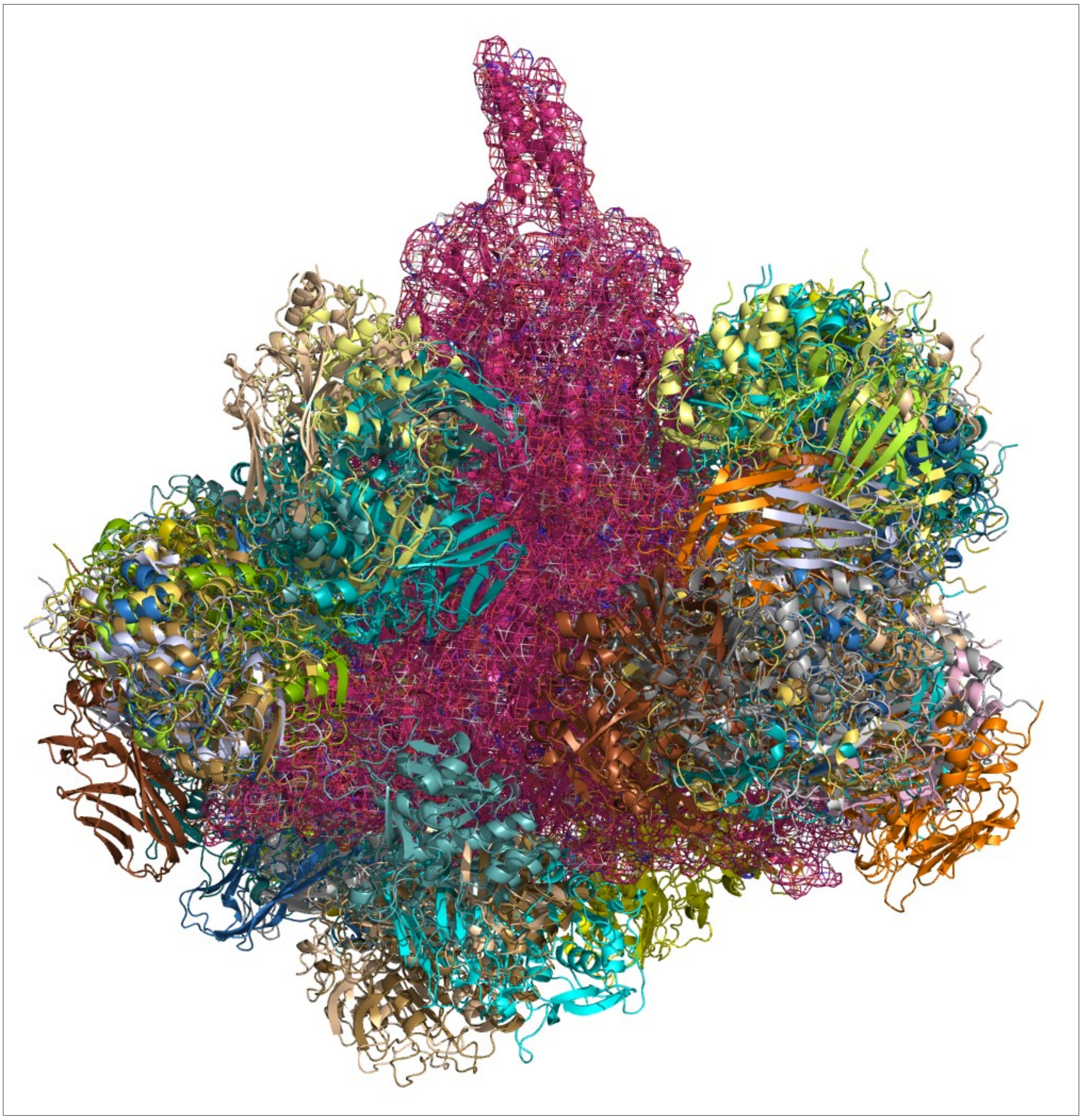

**Table S2. Scores for top 100 CoV-2 Spike–Furin docked poses ranked by $S_{dock}$** (see section **2.3.2.1 - 2.3.2.3, Materials and Methods**).

| **Model ID (based on rank)** | **$BSA_{PRRAR}$ (Å$^2$)** | **$nBSA_{PRRAR}$** | **$Sc^{FLCS}$** | **$S_{dock}$** |
|---|---|---|---|---|
| RR1 | 546.128 | 0.247 | 0.770 | 0.972 |
| RR2 | 467.328 | 0.241 | 0.696 | 0.906 |
| RR3 | 544.543 | 0.210 | 0.716 | 0.876 |
| RR4 | 568.184 | 0.216 | 0.691 | 0.864 |
| RR5 | 435.363 | 0.228 | 0.661 | 0.858 |
| RR6 | 457.662 | 0.238 | 0.631 | 0.852 |
| RR7 | 569.994 | 0.191 | 0.712 | 0.847 |
| RR8 | 391.570 | 0.189 | 0.711 | 0.843 |
| RR9 | 533.784 | 0.221 | 0.642 | 0.834 |
| RR10 | 517.731 | 0.146 | 0.755 | 0.832 |
| RR11 | 366.049 | 0.156 | 0.730 | 0.821 |
| RR12 | 384.993 | 0.150 | 0.733 | 0.816 |
| RR13 | 419.974 | 0.196 | 0.667 | 0.816 |
| RR14 | 384.423 | 0.195 | 0.668 | 0.816 |
| RR15 | 430.356 | 0.120 | 0.761 | 0.814 |
| RR16 | 540.292 | 0.198 | 0.657 | 0.811 |
| RR17 | 339.607 | 0.162 | 0.710 | 0.810 |
| RR18 | 203.347 | 0.076 | 0.784 | 0.805 |
| RR19 | 428.685 | 0.191 | 0.661 | 0.805 |
| RR20 | 354.425 | 0.138 | 0.727 | 0.799 |
| RR21 | 552.749 | 0.119 | 0.741 | 0.795 |
| RR22 | 406.678 | 0.106 | 0.752 | 0.794 |
| RR23 | 467.913 | 0.160 | 0.688 | 0.788 |
| RR24 | 143.555 | 0.066 | 0.771 | 0.787 |
| RR25 | 348.837 | 0.114 | 0.737 | 0.787 |
| RR26 | 475.195 | 0.189 | 0.637 | 0.782 |
| RR27 | 297.038 | 0.106 | 0.739 | 0.782 |
| RR28 | 480.299 | 0.190 | 0.635 | 0.782 |
| RR29 | 368.961 | 0.119 | 0.726 | 0.780 |
| RR30 | 319.929 | 0.102 | 0.739 | 0.778 |
| RR31 | 532.620 | 0.203 | 0.603 | 0.776 |
| RR32 | 152.871 | 0.059 | 0.755 | 0.768 |
| RR33 | 497.013 | 0.147 | 0.673 | 0.760 |

| | | | | |
|---|---|---|---|---|
| RR34 | 452.581 | 0.151 | 0.667 | 0.760 |
| RR35 | 415.054 | 0.141 | 0.678 | 0.759 |
| RR36 | 654.652 | 0.175 | 0.626 | 0.754 |
| RR37 | 327.674 | 0.068 | 0.734 | 0.752 |
| RR38 | 456.972 | 0.145 | 0.664 | 0.750 |
| RR39 | 335.308 | 0.142 | 0.668 | 0.750 |
| RR40 | 567.676 | 0.180 | 0.613 | 0.750 |
| RR41 | 524.557 | 0.199 | 0.575 | 0.749 |
| RR42 | 486.607 | 0.192 | 0.589 | 0.748 |
| RR43 | 221.766 | 0.067 | 0.728 | 0.746 |
| RR44 | 382.312 | 0.122 | 0.685 | 0.745 |
| RR45 | 498.099 | 0.224 | 0.515 | 0.745 |
| RR46 | 315.677 | 0.124 | 0.682 | 0.744 |
| RR47 | 389.157 | 0.096 | 0.706 | 0.743 |
| RR48 | 422.396 | 0.107 | 0.691 | 0.738 |
| RR49 | 479.041 | 0.237 | 0.467 | 0.736 |
| RR50 | 561.150 | 0.158 | 0.629 | 0.735 |
| RR51 | 384.495 | 0.137 | 0.651 | 0.730 |
| RR52 | 211.027 | 0.089 | 0.697 | 0.729 |
| RR53 | 213.287 | 0.065 | 0.710 | 0.727 |
| RR54 | 278.004 | 0.123 | 0.663 | 0.726 |
| RR55 | 556.757 | 0.169 | 0.599 | 0.724 |
| RR56 | 345.334 | 0.169 | 0.599 | 0.724 |
| RR57 | 324.116 | 0.132 | 0.646 | 0.719 |
| RR58 | 617.661 | 0.135 | 0.638 | 0.716 |
| RR59 | 241.915 | 0.071 | 0.692 | 0.713 |
| RR60 | 268.053 | 0.089 | 0.676 | 0.709 |
| RR61 | 312.872 | 0.097 | 0.668 | 0.707 |
| RR62 | 277.198 | 0.122 | 0.641 | 0.705 |
| RR63 | 378.888 | 0.136 | 0.624 | 0.704 |
| RR64 | 327.364 | 0.069 | 0.682 | 0.702 |
| RR65 | 398.884 | 0.151 | 0.598 | 0.700 |
| RR66 | 533.770 | 0.192 | 0.517 | 0.693 |
| RR67 | 338.996 | 0.111 | 0.639 | 0.693 |
| RR68 | 312.146 | 0.139 | 0.602 | 0.688 |
| RR69 | 411.373 | 0.131 | 0.608 | 0.685 |
| RR70 | 358.715 | 0.135 | 0.598 | 0.681 |

| | | | | |
|---|---|---|---|---|
| RR71 | 253.788 | 0.090 | 0.643 | 0.678 |
| RR72 | 404.504 | 0.102 | 0.628 | 0.674 |
| RR73 | 209.889 | 0.055 | 0.660 | 0.673 |
| RR74 | 282.161 | 0.094 | 0.631 | 0.670 |
| RR75 | 222.060 | 0.052 | 0.657 | 0.669 |
| RR76 | 279.145 | 0.066 | 0.648 | 0.667 |
| RR77 | 340.124 | 0.078 | 0.640 | 0.667 |
| RR78 | 237.750 | 0.056 | 0.652 | 0.666 |
| RR79 | 313.640 | 0.094 | 0.627 | 0.666 |
| RR80 | 273.857 | 0.061 | 0.648 | 0.664 |
| RR81 | 273.857 | 0.061 | 0.648 | 0.664 |
| RR82 | 217.383 | 0.055 | 0.649 | 0.662 |
| RR83 | 455.640 | 0.152 | 0.551 | 0.661 |
| RR84 | 237.896 | 0.067 | 0.638 | 0.658 |
| RR85 | 239.638 | 0.057 | 0.644 | 0.658 |
| RR86 | 207.618 | 0.053 | 0.645 | 0.657 |
| RR87 | 410.788 | 0.096 | 0.613 | 0.655 |
| RR88 | 310.860 | 0.086 | 0.620 | 0.654 |
| RR89 | 326.202 | 0.085 | 0.619 | 0.652 |
| RR90 | 294.880 | 0.081 | 0.618 | 0.648 |
| RR91 | 241.056 | 0.051 | 0.627 | 0.639 |
| RR92 | 313.072 | 0.082 | 0.606 | 0.637 |
| RR93 | 309.123 | 0.099 | 0.590 | 0.636 |
| RR94 | 352.503 | 0.105 | 0.584 | 0.636 |
| RR95 | 394.385 | 0.111 | 0.571 | 0.631 |
| RR96 | 266.771 | 0.068 | 0.603 | 0.625 |
| RR97 | 325.878 | 0.151 | 0.508 | 0.624 |
| RR98 | 378.979 | 0.097 | 0.573 | 0.619 |
| RR99 | 309.579 | 0.082 | 0.584 | 0.616 |
| RR100 | 309.579 | 0.082 | 0.584 | 0.616 |

**Table S3. Furin anionic residues (Asp, Glu) having the potential to form interfacial salt-bridges with FLCS$_{Spike}$.** Exposed (or partially exposed) anionic residues proximal to the Furin catalytic triad are highlighted with a background color 'light yellow'.

| **Residues** | **D$_{triad}$** | ***bur*** | **Burial status** |
|---|---|---|---|
| 153-ASP | 3.264 | 0.00 | buried |
| 162-ASP | 17.999 | 0.00 | buried |
| 157-GLU | 14.000 | 0.01 | buried |
| 201-GLU | 9.627 | 0.01 | buried |
| 331-GLU | 15.068 | 0.01 | buried |
| 154-ASP | 5.707 | 0.04 | buried |
| 174-ASP | 15.288 | 0.04 | buried |
| 168-ASP | 19.224 | 0.06 | partially exposed |
| 236-GLU | 14.153 | 0.08 | partially exposed |
| 301-ASP | 16.911 | 0.08 | partially exposed |
| 306-ASP | 14.002 | 0.08 | partially exposed |
| 355-ASP | 11.346 | 0.12 | partially exposed |
| 228-ASP | 11.503 | 0.15 | partially exposed |
| 233-ASP | 19.726 | 0.24 | partially exposed |
| 259-ASP | 17.834 | 0.26 | partially exposed |
| 299-GLU | 18.572 | 0.28 | partially exposed |
| 179-ASP | 19.908 | 0.31 | exposed |
| 258-ASP | 11.858 | 0.31 | exposed |
| 181-ASP | 18.167 | 0.37 | exposed |
| 362-GLU | 18.798 | 0.42 | exposed |
| 264-ASP | 17.833 | 0.46 | exposed |
| 191-ASP | 11.772 | 0.56 | exposed |
| 177-ASP | 19.117 | 0.57 | exposed |
| 257-GLU | 14.321 | 0.64 | exposed |
| 230-GLU | 18.719 | 0.72 | exposed |

**Table S4. Occurrence and average contact intensities of all unique salt-bridges at the SARS-CoV-2 Spike–Furin interface computed on the static ensemble of the top 100 (re-)ranked docked poses (as enlisted in Table S2).** '-S' & '-F' in the salt-bridge descriptor strings refer to the receptor and the ligand chains respectively. Rows corresponding to the arginine – salt-bridges falling within the pentapeptide $_{681}$PRRAR$_{685}$ motif (activation loop of FLCS$_{Spike}$) is highlighted in three different font colors for R682 (red), R683 (purple), R685 (green).

| ***Salt-bridge*** | ***TotC*** | ***Frames$_p$*** | ***ACI*** | ***Occ*** |
|---|---|---|---|---|
| *214-ARG-S↔112-GLU-F* | *2* | *1* | *2.00* | *0.01* |
| *654-GLU-S↔357-ARG-F* | *4* | *1* | *4.00* | *0.01* |
| *654-GLU-S↔193-ARG-F* | *6* | *2* | *3.00* | *0.02* |
| *682-ARG-S↔233-ASP-F* | *7* | *3* | *2.33* | *0.03* |
| *683-ARG-S↔230-GLU-F* | *5* | *3* | *1.67* | *0.03* |
| *685-ARG-S↔306-ASP-F* | *11* | *6* | *1.83* | *0.06* |
| *682-ARG-S↔236-GLU-F* | *26* | *11* | *2.36* | *0.11* |
| *685-ARG-S↔236-GLU-F* | *46* | *17* | *2.71* | *0.17* |
| *682-ARG-S↔230-GLU-F* | *52* | *18* | *2.89* | *0.18* |
| *683-ARG-S↔236-GLU-F* | *57* | *25* | *2.28* | *0.25* |
| *683-ARG-S↔264-ASP-F* | *64* | *28* | *2.29* | *0.28* |
| *685-ARG-S↔264-ASP-F* | *86* | *31* | *2.77* | *0.31* |

***TotC:*** *Total Counts (Ion-pairs)*
***Frames$_p$:*** *Number of frames the salt-bridge (Residue-Pair) is found in*
***ACI:*** *Average Contact Intensity = TotC/Frames$_p$*
***Occ:*** *Occurrence = Frames$_p$/Frames$_t$*
*Frames$_t$=Total number of models in the static ensemble: 100*

**Figure S4. The static Spike – Furin ensemble comprising of the plausible (top 50 re-ranked) Furin poses docked onto Spike with the selected top ranked (RR1$_{CoV-2}$) docked pose highlighted.**

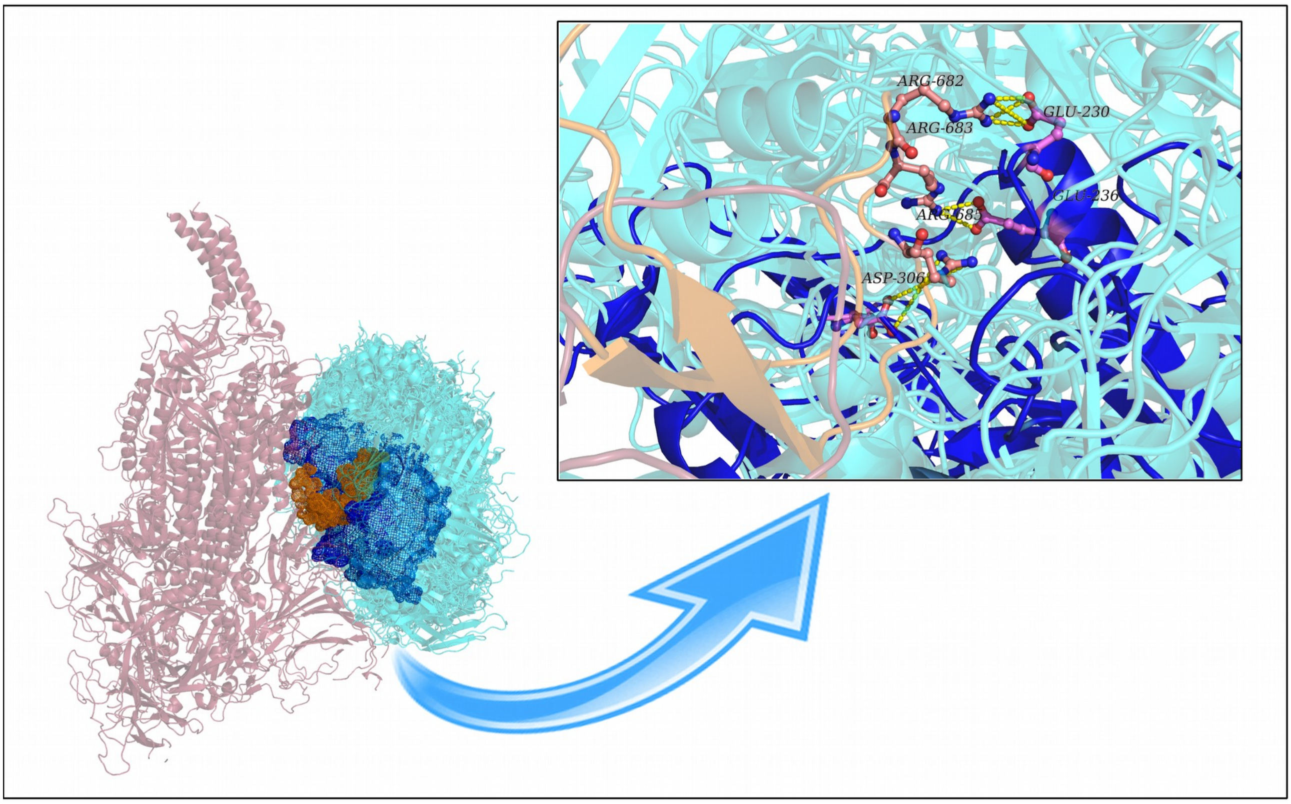

**Video S1. The Spike–Furin binding in SARS-CoV-2: Essential Dynamics of Interfacial Salt-bridges portrayed** (see '*unlisted*' YouTube link: [https://www.youtube.com/watch?v=wsHKpr9gZ9E](https://www.youtube.com/watch?v=wsHKpr9gZ9E)). Video built in PyMol based on a 10 ns epoch (i.e., 1000 frames, 150-160 ns) extracted from the 300 ns long trajectory of $RR1_{CoV2}$ portraying the high persistence salt-bridges at the Spike–Furin interface in real time. The catalytic triad is shown in dots.

***Table S5. Persistence and average contact intensities of all unique salt-bridges at the SARS-CoV-2 Spike–Furin interface for ZR1$_{CoV\text{-}2}$ (zdock + IraPPA re-ranking) along its 100 ns MD simulation trajectory.*** *'-S' & '-F' in the salt-bridge descriptor strings refer to the receptor and the ligand chains respectively. Rows corresponding to the arginine – salt-bri*dges falling within the pentapeptide $_{681}$PRRAR$_{685}$ motif (activation loop of FLCS$_{Spike}$) is highlighted in three different font colors for R682 (red), R683 (purple), R685 (green).

| *Salt-bridge* | *TotC* | *Frames$_p$* | *ACI* | *Pers* |
|---|---|---|---|---|
| *683-ARG-S↔258-ASP-F* | *1* | *1* | *1.00* | *0.00010* |
| *811-LYS-S↔460-ASP-F* | *2* | *2* | *1.00* | *0.00020* |
| *685-ARG-S↔230-GLU-F* | *5* | *5* | *1.00* | *0.00050* |
| *627-ASP-S↔197-ARG-F* | *8* | *8* | *1.00* | *0.00080* |
| *825-LYS-S↔500-ASP-F* | *15* | *13* | *1.15* | *0.00130* |
| *214-ARG-S↔112-GLU-F* | *22* | *18* | *1.22* | *0.00180* |
| *683-ARG-S↔236-GLU-F* | *47* | *46* | *1.02* | *0.00460* |
| *936-ASP-S↔497-ARG-F* | *130* | *83* | *1.57* | *0.00830* |
| *214-ARG-S↔136-GLU-F* | *579* | *337* | *1.72* | *0.03370* |
| *683-ARG-S↔230-GLU-F* | *524* | *405* | *1.29* | *0.04050* |
| *685-ARG-S↔264-ASP-F* | *886* | *563* | *1.57* | *0.05631* |
| *654-GLU-S↔193-ARG-F* | *962* | *567* | *1.70* | *0.05671* |
| *627-ASP-S↔193-ARG-F* | *809* | *587* | *1.38* | *0.05871* |
| *627-ASP-S↔357-ARG-F* | *3975* | *1362* | *2.92* | *0.13621* |
| *214-ARG-S↔131-ASP-F* | *4315* | *2041* | *2.11* | *0.20412* |
| *683-ARG-S↔264-ASP-F* | *4649* | *2740* | *1.70* | *0.27403* |
| *627-ASP-S↔359-LYS-F* | *4029* | *3004* | *1.34* | *0.30043* |
| *683-ARG-S↔259-ASP-F* | *9022* | *4891* | *1.84* | *0.48915* |
| *685-ARG-S↔236-GLU-F* | *33470* | *9689* | *3.45* | *0.96900* |
| *682-ARG-S↔230-GLU-F* | *38134* | *9911* | *3.85* | *0.99120* |

***TotC:*** *Total Counts (Ion-pairs)*
***Frames$_p$:*** *Number of frames the salt-bridge (Residue-Pair) is found in*
***ACI:*** *Average Contact Intensity = TotC/ Frames$_p$*
***Pers:*** *Persistence = Frames$_p$/ Frames$_t$*
*Frames$_t$ = Total number of frames=10000 (sampled at 10 ps interval)*

**Figure S5. Frequency distributions of salt-bridge persistence and ACIs in RR1$_{CoV-2}$ and ZR1$_{CoV-2}$**

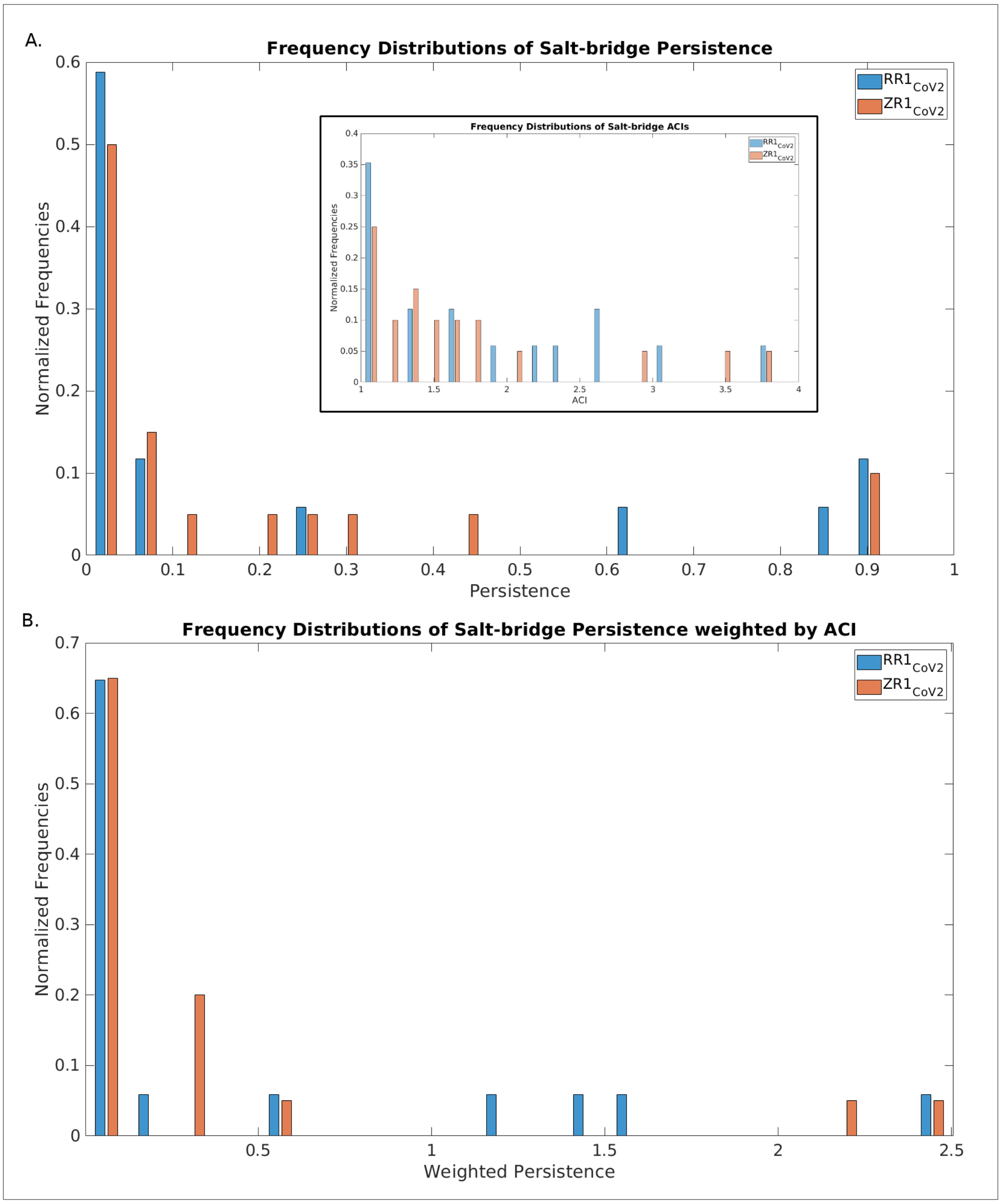

**Table S6. Persistence and average contact intensities of all unique salt-bridges at the SARS-CoV Spike–Furin interface for ZR1$_{CoV}$ *(zdock+ IRAppA re-ranking)* along its 100 ns MD simulation trajectory.** '-S' & '-F' in the salt-bridge descriptor strings refer to the receptor and the ligand chains respectively. Rows corresponding to high persistence interfacial salt-bridges are highlighted in different font colors unique to their cationic partners (indigo: R667, dark green: K672) coming from the SARS-CoV FLCS$_{Spike}$.

| Salt-bridge | TotC | *Frames$_p$* | ACI | Pers |
|---|---|---|---|---|
| 208-ASP-S↔349-LYS-F | 13 | 13 | 1.00 | 0.00130 |
| 294-GLU-S↔298-ARG-F | 1 | 1 | 1.00 | 0.00010 |
| 640-GLU-S↔193-ARG-F | 25 | 25 | 1.00 | 0.00250 |
| 207-ARG-S↔131-ASP-F | 9 | 9 | 1.00 | 0.00090 |
| 208-ASP-S↔130-ARG-F | 765 | 231 | 3.31 | 0.02310 |
| 667-ARG-S↔258-ASP-F | 2865 | 2787 | 1.03 | 0.27870 |
| 672-LYS-S↔258-ASP-F | 9167 | 5839 | 1.57 | 0.58390 |
| 667-ARG-S↔230-GLU-F | 11971 | 6147 | 1.95 | 0.61470 |
| 667-ARG-S↔257-GLU-F | 36176 | 9505 | 3.81 | 0.95050 |

***TotC:*** *Total Counts (Ion-pairs)*
***Frames$_p$:*** *Number of frames the salt-bridge (Residue-Pair) is found in*
***ACI:*** *Average Contact Intensity = TotC/ Frames$_p$*
***Pers:*** *Persistence = Frames$_p$/ Frames$_t$*
*Frames$_p$ = Total number of frames=10000 (sampled at 10 ps interval)*

**Figure S6. Interfacial ionic bond network in ZR1$_{\text{CoV}}$ (the baseline) shows the proximal looping of the FLCS$_{\text{Spike}}$ enabling R667 and K672 to be involved in multiple persistent as well as interchangeable Spike – Furin interfacial salt-bridges in SARS-CoV.**

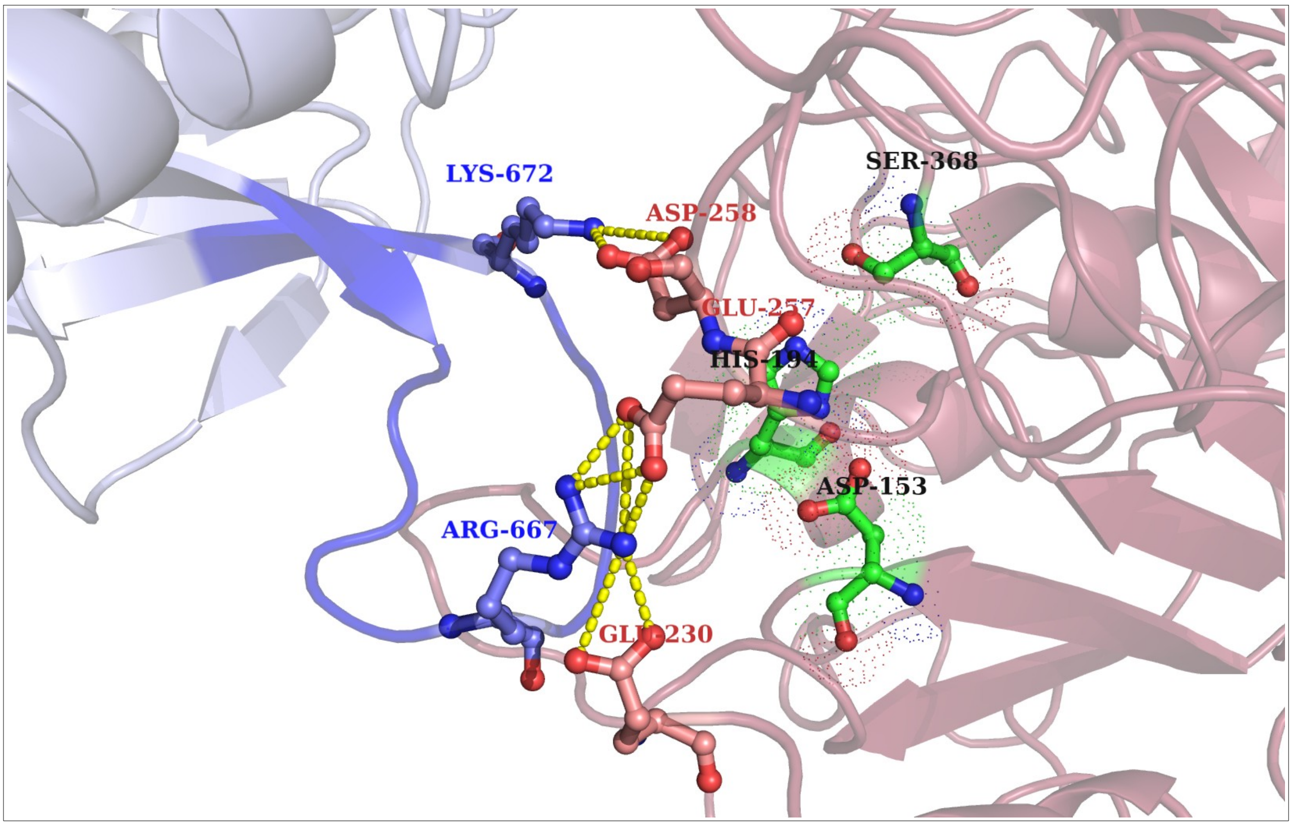

**Figure S7. Entropy arrest in the Spike–Furin interactions (SARS-CoV-2, SARS-CoV) as revealed in $RR1_{CoV\text{-}2}$, $ZR1_{CoV}$**

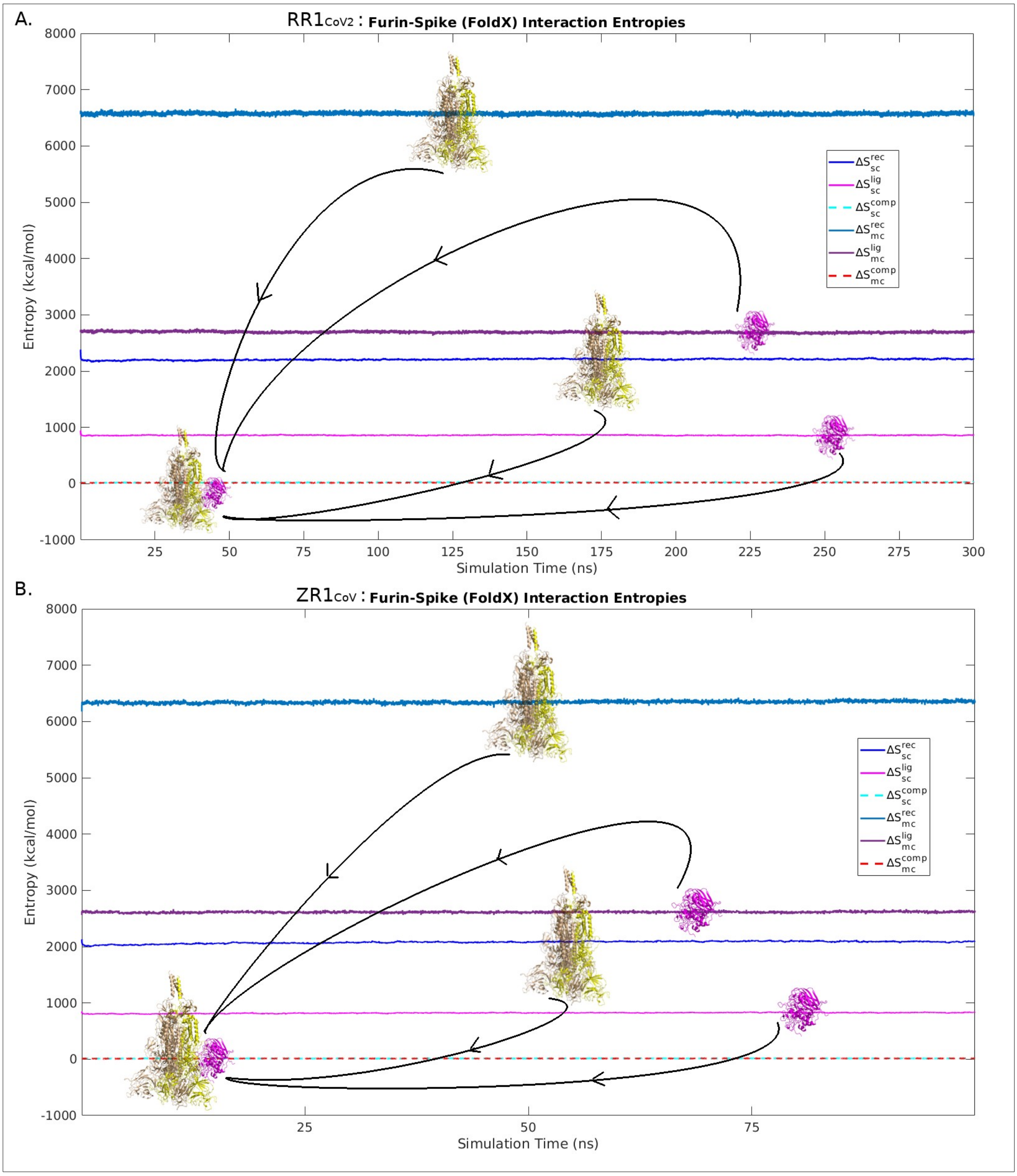

**Figure S8. Overlaid Ramachrandran Plots for FLCS$_{Spike}$ pertaining to (A) unbound and (B) Furin-bound Spike states.** Each plot is overlaid with 100 atomic models belonging to the same state (unbound or bound). Within each individual plot (i.e., pertaining to each atomic model), the contiguity of {Φ, Ψ} points for successively connected residues in the FLCS$_{Spike}$ loop is portrayed by adding the successively connected residues by thin dashed black lines (--) belonging to the -P$_{681}$-R$_{682}$-R$_{683}$-A$_{684}$-R$_{685}$- pentapeptide motif.

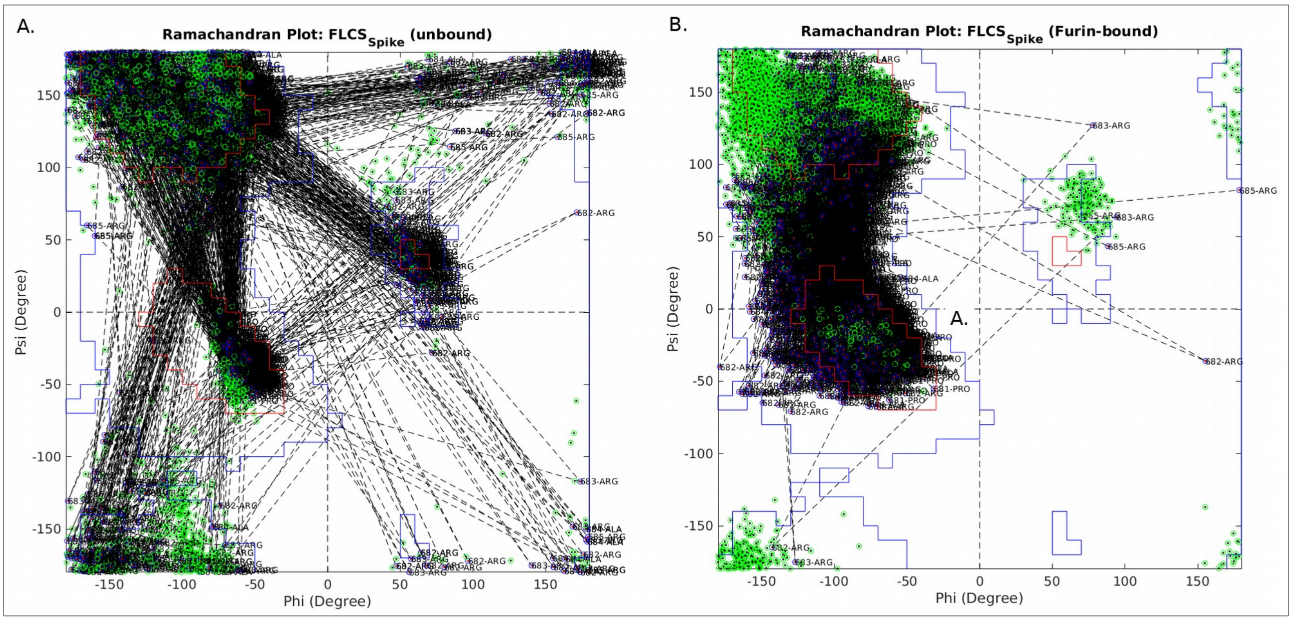

***References***


1. Edgar, R.C. MUSCLE: A Multiple Sequence Alignment Method with Reduced Time and Space Complexity. *BMC Bioinformatics* **2004**, *5*, 113, doi:10.1186/1471-2105-5-113.
2. Sievers, F.; Higgins, D.G. Clustal Omega for Making Accurate Alignments of Many Protein Sequences: Clustal Omega for Many Protein Sequences. *Protein Sci.* **2018**, *27*, 135–145, doi:10.1002/pro.3290.
